# Tethering-mediated recruitment of pioneer factors by NuRD enables signal dependent gene activation and protects cell identity

**DOI:** 10.64898/2026.08.11.744124

**Authors:** Rita S. Monteiro, Annika Charlotte Sell, Kristina B. Emdal, Sarah L. Lundregan, Christina M. Schuh, Silvia Raineri, François Dossin, Edith Heard, Jesper V. Olsen, Joshua M. Brickman

## Abstract

Cell identity relies on precise control of transcription factor (TF) activity, localization and interpretation of signaling. Mitogen-activated protein kinase (MAPK) signaling regulates transcription via the ERK kinase, phosphorylating components of the Mediator complex to regulate its association with TF-bound enhancers. Here we describe two distinct forms of TF association related to signaling response, stable DNA recognition and tethering to active sites of transcription. We find that the tethering of TFs at these active sites is both essential for signaling response and cell identity. To identify tethering substrates, we focused on signaling dependent associations with the transcriptional coregulator, the Mediator, and find the NuRD complex is engaged with the Mediator at active promoters and enhancers. Here, NuRD functions as a molecular scaffold recruiting pioneer TFs like SOX2 independently of DNA sequence, enhancing ERK dependent transcription while sequestering these factors away from context inappropriate binding sites elsewhere in the genome. Balancing the recognition of a restricted spectrum of canonical DNA sites by pioneer factors while retaining them at high concentrations in the vicinity of transcription supports robust transcriptional response, ensures plasticity in differentiation and explains why these factors bind a fraction of their naturally occurring sites in any given cell type.

## Introduction

Cell identity is largely defined by the genes that are actively transcribed, giving transcriptional regulation a central role in cell fate determination. Transcriptional control involves multiple steps and molecular components, including the integration of signaling pathways that converge in the nucleus. There, specialized proteins known as transcription factors (TFs) bind DNA to directly regulate transcription. Despite the sixty years since the discovery of the first TF^1^, it remains a mystery how gene-specific regulation by TFs is achieved. While DNA in cells is wrapped in chromatin, certain TFs have the capacity to recognize their sites in chromatin, yet despite this ability these pioneer factors only bind a fraction of their recognition sites in a given cell type^2^.

Transcription by RNA Polymerase II (POLII) is regulated by a combination of signal transduction and sequence specific DNA binding domain proteins or TFs^3,4^. These TFs generally recognize two sets of regulatory elements: promoters, located proximal to the transcription start site (TSS) and distal enhancers. TFs are thought to recognize specific sequences through their DNA binding domain and use distinct regulatory domains to recruit cofactors and stimulate transcription^5,6^. One of the major cofactors thought to bridge the interface between enhancer bound TFs and POLII is the Mediator complex, a large multiprotein complex, that is known to associate with the C-terminal tail of POLII, playing a vital role in transmitting signals from the TFs to the polymerase^7–9^.

Development and differentiation exploit the coupling of transcriptional regulation to signal transduction to regulate changes in cell states. Early embryonic development and mouse embryonic stem cells (mESCs) represent an excellent paradigm to explore the signaling harness TF activity to regulate the dynamics of cell state change^10^. mESCs are derived based on the *ex vivo* expansion of the inner cell mass (ICM) of the mammalian blastocyst^11,12^. mESCs are pluripotent, as they can differentiate into all embryonic lineages and their differentiation depends on kinase signaling via the FGF/ERK pathway^13–18^. Intracellular signaling via this pathway, known as mitogen activated protein kinase (MAPK) pathway, consists of the three consecutive protein kinases RAF, MEK and ERK, which are activated by sequentially phosphorylating each other^19^ and culminates in phospho-ERK translocation into the nucleus to regulate transcription^20^.

In mESCs, ERK acts on a network of TFs known as the pluripotency network that includes factors such as NANOG, OCT4, SOX2, KLF4 and ESRRB^21^. In the nucleus ERK does not act directly on the pluripotency network but phosphorylates a variety of transcriptional coactivators to regulate enhancer activity, including the tail subunit 24 of the Mediator complex (MED24)^18^. The phosphorylation of MED24 and associated cofactors results in the decommissioning of enhancers associated with pluripotency and the commissioning of those related to differentiation. ERK dependent enhancer decommissioning results in the disassociation of POLII and associated factors but leaves the pluripotency-associated TFs at these decommissioned enhancers, where they maintain a plastic state and block commitment. Although pluripotency TFs do not leave decommissioned enhancers, we observed migration of these TFs to sites of active transcription that lack their consensus motifs^18^. Thus, there appear two types of TF binding; high affinity site specific interactions that are stable through transcriptional change and low affinity, tethering-mediated recruitment to the transcription complex.

A subset of TFs are agnostic to the chromatin state of their target sites and have been described as pioneer factors ^22,23^. The rewiring of transcriptional programs in response to signal transduction occurs as these TFs are constitutively present in the nucleus and have thousands of recognizable DNA binding motifs that they do not recognize. What determines the set of pioneer sites recognized in one cell type and prevents them from eroding cell identity by recognizing inappropriate sites in any given cell type? To address this question and the relevance of TF tethering to supporting plasticity and protecting cell identity, we focused on colocalization of TFs and coactivators of MED24 response to signaling. We found that both pluripotency pioneer factors, SOX2 and ESRRB, were recruited via tethering to active sites of transcription enriched with the AP1 and ATF3 DNA footprints. Using Rapid Immunoprecipitation Mass spectrometry of Endogenous proteins (RIME)^24^ we identify mediators of tethering via MED24, and identified multiple components of the nucleosome remodeling and deacetylation (NuRD) complex. An abundant, highly conserved multiprotein complex, that was initially defined as a transcriptional corepressor^25–27^, we found that this complex was recruited alongside the Mediator and TFs as part of the signaling dependent transcriptional activation responses. Recruitment of NuRD to sites of activation was necessary for the tethering of TFs at sites of transcriptional activity, both enhancers and promoters. This recruitment was necessary for the activation of signaling dependent transcription, and in its absence, pioneer factors like SOX2 bind throughout the genome to deconstruct cellular identity. Taken together the NuRD complex retains effective TF levels in the vicinity of active transcription where it supports robust transcriptional response and retains a pool of TFs that ensures flexibility in differentiation, while maintaining a limiting pioneer TF concentration elsewhere to ensure transcription is restricted to the correct genomic sites.

## Results

### Tethering-mediated recruitment of pioneer factors to Mediator-bound ERK-induced transcription sites

To identify factors associated with signaling mediated regions of transcriptional change we focused on the Mediator complex. We previously identified MED24 – a Mediator tail subunit known to interact with TFs and co-factors - as a crucial regulator of the ERK-induced transcriptional response in mouse embryonic stem cells (mESCs). To discern signaling-dependent associations with the Mediator complex, we exploited a cell line, previously described, that includes a cRAF-ER^T2^ (*Z*)-4-hydrotamoxifen (4OHT)-inducible fusion protein contained in an *Med24* KO mESC line with an rtTA inducible FLAG-tagged MED24. We re-analyzed the binding of the FLAG-tagged MED24 and characterized the recruitment of MED24 after 2h of ERK induction (Fig. 1A; Table S1A-C, see methods for details on ERK activation). We compared the genomic regions that gain, lose or maintain the binding of MED24 to MED1, P300, H3K27ac and the pluripotency pioneer TFs; ESRRB and SOX2^18^ (Fig. 1B). Analysis of the DNA binding motifs present at these MED24 sites confirmed that these represent ERK regulatory elements in mESCs – JUN/AP1 and ATF3 motifs are enriched in sites that gain MED24 binding and motifs of pluripotency factors like ESRRB, SOX2 and POU-SOX-NANOG-TCF at regions that lose it upon ERK activation (Fig. 1C; Table S1D-F). These observations suggest that pioneer factors such as SOX2 and ESRRB, remain stably associated with pluripotency TFs consensus sites, but are also recruited to sites of active transcription, moving in response to signaling. Differential ATAC-seq footprint at SOX2 and ESRRB peaks confirms that new binding in response to signaling occurs in the absence of the SOX or ESRRB footprint on DNA with these TFs both being recruited to regions footprinted by JUN/AP1 and ATF3 (Fig. 1D). Moreover, these TFs move away from sites where they also do not directly appear to recognize DNA, but rather to sites containing motifs for other members of the pluripotency network. This confirmed that MED24 and the Mediator complex moved rapidly in response to ERK signaling and that pluripotency pioneer TFs both remain stably associated with high affinity sites, but move to and from sites where they appear to be tethered by other TFs.

**Figure 1.**
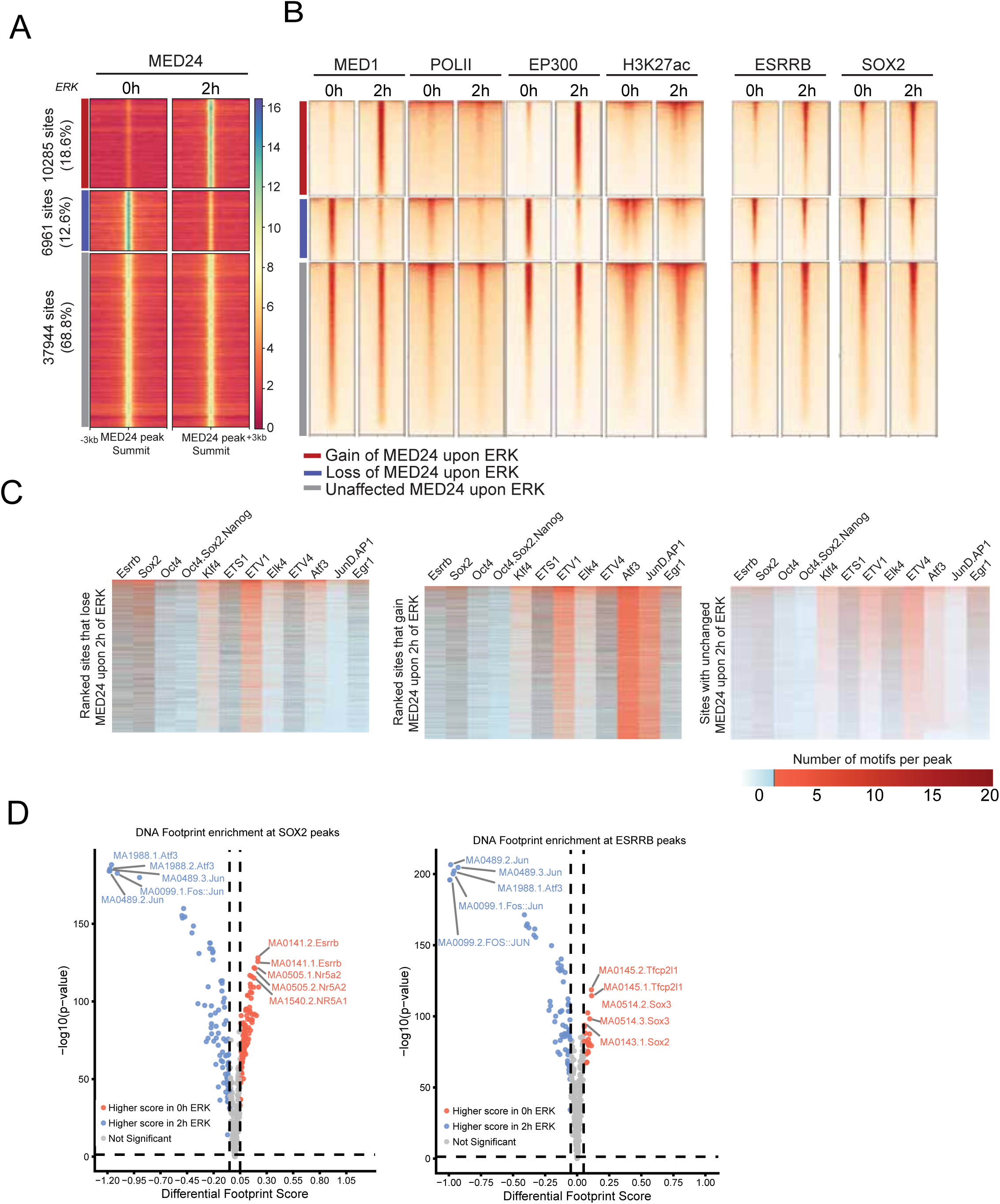
Mediator and transcriptional machinery movement in response to ERK is followed by tethering-mediated recruitment of pioneer factors. (**A**) Heatmap of hierarchical binding of MED24 at genomic regions with unchanged binding of MED24 (grey) and regions identified as differentially bound, either increase in binding upon ERK (red) or loss of binding upon 2h of ERK activity (blue). Genomic coordinates are in Tables S1A-C. (**B**) Heatmap of hierarchical binding in the absence of and upon 2h of ERK activation of MED24, MED1, POLII, EP300, H3K27ac, ESRRB and SOX2 at genomic regions with either unchanged, gain or loss of MED24 binding upon ERK. Co-factor and TF binding data from Hamilton et al., 2019. (**C**) Peaks ranked by MED24 loss or gain upon ERK activation and density of known DNA-binding motifs at each peak. Sites that lose MED24 have more motifs for pluripotency TFs like ESRRB, SOX2 and KLF4; while sites that gain MED24 are enriched with ATF3 and JUND (AP1) motifs (Table S1D-F). (**D**) Volcano plot depicting differential footprint scores and significance (-log10(pvalue), calculated by TOBIAS) for DNA binding motifs found at SOX2 (left) and ESRRB (right) peaks in response to ERK signaling. The differential score was calculated on consensus ESRRB and SOX2 peaks from 0h and 2h ERK ChIP experiments from Hamilton et al., 2019 and accessibility differences based on ATAC signals from the same conditions.

### The NuRD complex is recruited with the Mediator to active sites of transcription in response to ERK signaling

The strong link between ERK and MED24 makes it the ideal proxy to investigate tethering substrates of the ERK transcriptional response. Using the FLAG-tagged MED24 we performed Rapid Immunoprecipitation Mass spectrometry of Endogenous proteins (RIME)^24^ to identify MED24 interactors in the presence and absence of ERK signaling in two independent cell clones (Fig. S1A). In the RIME, we identified and quantified 1305 proteins with an overlap of 1031 proteins between clones (Table S2 and Fig. S1B). In addition, we analyzed the nuclear proteomes, and we identified and quantified in total 5994 proteins (Fig. S1C, Table S3), respectively. The nuclear proteome confirmed increased MED24 abundance for DOX-induced conditions and active ERK signaling with increased protein abundance of immediate early genes (IEGs) such as EGR1, ARC and c-JUN upon induction of the cRAF-ERT2 construct (Fig. S1D). The MED24-induced RIME experiment showed specific enrichment of Mediator subunits by correlation with nuclear proteome abundance for MED24-induced conditions (Fig. S1E and F). This supported the validity of the RIME to further explore the endogenous interaction partners of MED24. We assessed the dynamic recruitment of MED24 associated proteins upon active ERK signaling (Fig. 2A and Fig. S1G) and identified 95 proteins that showed increased interaction with MED24 following 2h of ERK induction in two independent cell lines, including 37 proteins that are also directly phosphorylated by ERK (Fig.2B and Fig.S1H). Consistent with the tethering of ESRRB observed in Fig. 1, we observe recruitment of ESRRB and other pioneer TFs associated with pluripotency associated with the ERK activated mediator complex (Table S2).

**Figure 2.**
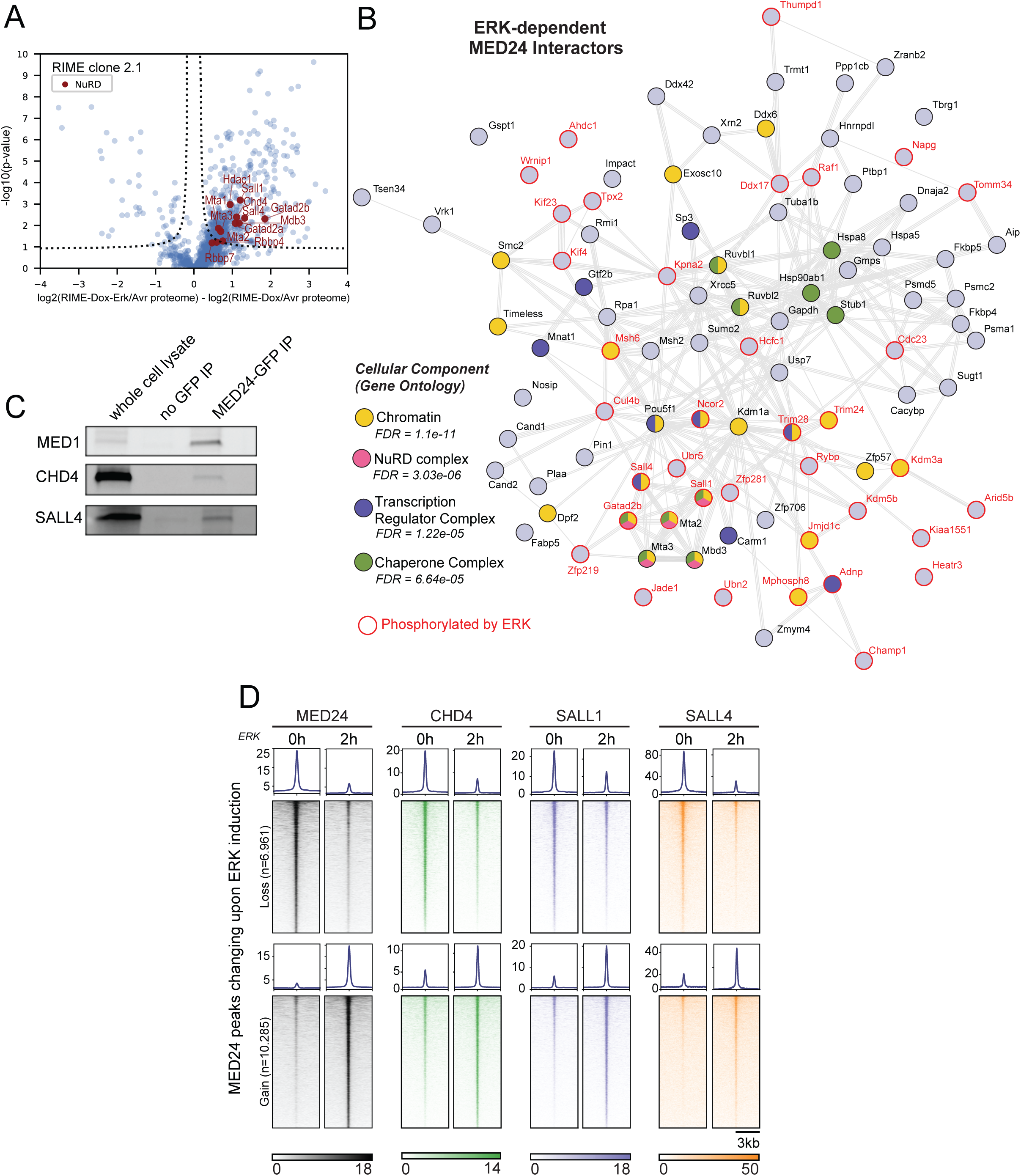
The NuRD complex is a an ERK-dependent interactor of MED24. **(A)** Volcano plot of clone 2.1 RIME data highlighting NuRD complex-related proteins (red). - log10(p-value) is plotted against log2 fold-change for the ERK-induced condition versus control. **(B)** Protein-protein interaction network of the 95 ERK-dependent MED24 interactors by gene name. Proteins with red circle are phosphorylated by ERK (Hamilton et al., 2019) and proteins belonging to the most relevant GO-terms are coloured according to the legend. **(C)** Western blot of IP pulldown of MED24-GFP, showing the native interaction with CHD4, a major subunit of the NuRD complex and SALL4, a co-factor of the complex. **(E)** Heatmap of the binding of MED24, CHD4, SALL1 and SALL4 at genomic regions that gain and lose binding of MED24 (as shown in Figure 1A) upon 2h of ERK activity (0h represents cells without ERK signaling, treated with PD17).

While the set of signaling-enhanced MED24 interactors included proteins related to chromatin, transcription regulator complex and chaperone complex, the most prominent class in this data were members of the NuRD complex. NuRD has two distinct enzymatic activities, executed by CHD4, a SWI/SNF-type ATPase for repositioning of nucleosomes and HDAC1/2 for lysine residue deacetylation. The complex also contains two MTA proteins (MTA1/2 and/or 3), histone chaperones RBBP4 and RBBP7, either GATAD2a or GATAD2b, CDK2AP1, and methyl-CpG-binding protein MBD2 or MBD3^28,29^. GATAD2b, MTA2, MTA3, and MBD3 as well as NuRD-associated TFs SALL1 and SALL4. Four out of these six proteins are directly phosphorylated by ERK (Fig. 2B, Fig. S1I). While not all NuRD components are enriched in response to signaling in both clones, Fig. S1I shows all are associated with the Mediator itself. Despite NuRD being predominantly associated with transcriptional repression, it has previously been identified as an interactor of the Mediator complex in neural stem cells^30^.

Since the original FLAG-MED24 cell-line was based on a transgene expression, we created an endogenously tagged MED24-mAID-GFP cell line that we used to confirm the native association of endogenous MED24 with components of the NuRD complex, CHD4 and SALL4 (Fig.2C). This interaction was further assessed by Cleavage Under Targets and Release Using Nuclease (CUT&RUN)^31^. CHD4, SALL1 and SALL4 shifted their binding after 2 hours of ERK stimulation, colocalizing with MED24 on the chromatin following the Mediator complex to active sites of transcription, like SOX2 and ESRRB (Fig. 2D). Taken together the NuRD complex moves in response to ERK signaling and colocalizes on chromatin with the Mediator.

### CHD4 mediates the binding preferences of the pioneer factor SOX2

To determine the roles of NuRD in mediating pioneer factor binding in response to signaling we introduced a protein degradation allele of the ATP-dependent histone remodeling subunit and central NuRD scaffold, CHD4 (Fig. 3A, S2A) to an ERK-inducible cell line stably expressing the cell intrinsically inducible cRAF-ERT2 construct in the *tigre* locus (Fig. S2A and B). This cell line allowed us to assess the direct effect of the loss of NuRD upon signaling mediated transcription and TF association by rapidly degrading CHD4 after 2h of adding auxin and homogenously inducing ERK signaling with the addition of 4OHT.

**Figure 3.**
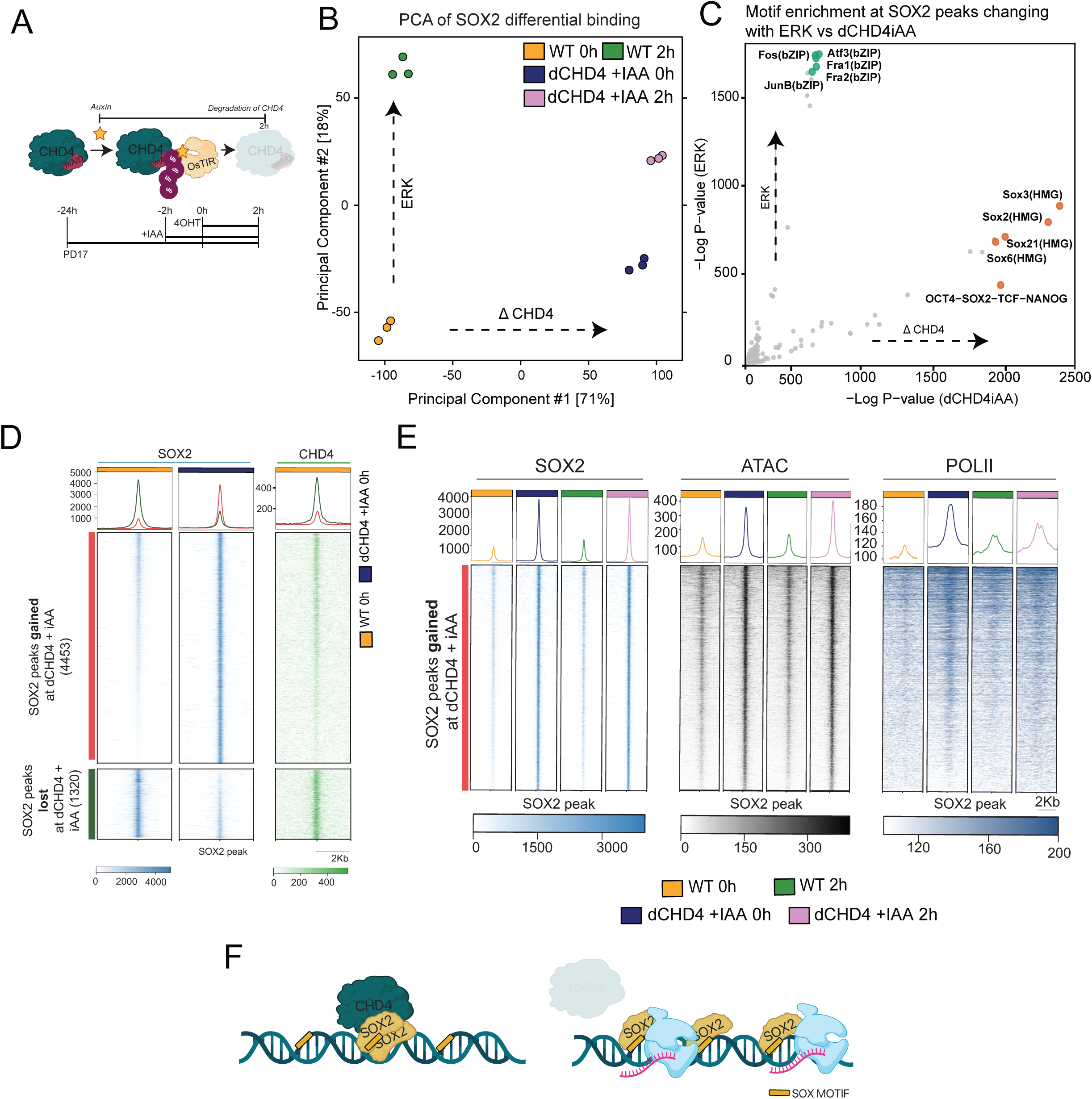
Loss of the NuRD complex leads to cell type inappropriate SOX2 binding and PolII recruitment. **(A)** Schematic showing the auxin-inducible degradation of CHD4 and experimental setup to access nascent transcription with TT_chem_-Seq upon ERK activation. (**B**) Principal component analysis (PCA) of the SOX2 binding in control and auxin-treated dCHD4 cells, with and without 2h of ERK activation. PC1 explains 71% of the total variance of the samples and separates samples by loss of CHD4, while a smaller variance in PC2 (18%) correlates with ERK activation. (**C**) Scatter plot comparing the significance of the enrichment of known DNA binding motifs found at SOX2 peaks with significant binding increase as a result of loss of CHD4 or ERK activation. Values were calculated using HOMER. (**D**) Heatmap of hierarchical binding of SOX2 and CHD4 at genomic regions with gain of SOX2 binding upon loss of CHD4 (red) and regions that lose SOX2 binding under the same conditions (green). Sites that lose SOX2 upon degradation of CHD4 show higher binding of the NuRD component that sites that gain SOX2.(**E**) Average profiles (top) and heatmaps (bottom) showing SOX2, ATAC-seq, and POLII signals centered on SOX2 peaks gained after CHD4 degradation. (**F**) Schematic summarizing the effect of the loss CHD4 on the binding of SOX2 to chromatin.

Of the pluripotent pioneer factors that move in response to ERK signaling, SOX2 has been shown in other contexts to interact with NuRD, and particularly CHD4^32,33^. We also observed that both before and after signaling, SOX2 occupies many of the same genomic regions as CHD4, with substantial overlap between their peaks (Fig. S2D; Table S4B,C). To investigate whether NuRD could regulate the recruitment of pioneer factors we examined SOX2 binding in the context of CHD4 loss and ERK activation. Differential analysis of SOX2 binding revealed that CHD4 depletion exerts a stronger effect on SOX2 occupancy than ERK induction (Fig. 3B; Table S5A-C), accounting for 71% of variance (dCHD4 + iAA vs WT), compared to signaling that is responsible for 18% (0h vs 2h). Upon CHD4 loss, 46% of SOX2 binding events were altered, with 35% representing novel binding in dCHD4 cells treated with iAA and cultured in PD17 (Table S5). These new SOX2-bound sites were enriched for canonical SOX motifs, suggesting direct and specific DNA binding (Fig. 3C). This contrasts to the influence of signaling, as described before, where SOX2 leaves non-sequence specific locations for sites of newly activated transcription enriched with ATF3/FOS motifs (Fig. 3C; Table S5D,E). While the new SOX2 binding events showed lower CHD4 occupancy prior to its degradation the sites that lost SOX2 upon CHD4 depletion were occupied by CHD4, (Fig. 3D; Table S5A,B), suggesting that CHD4 restrains SOX2 to specific genomic regions.

Given SOX2’s established role as a pioneer factor that promotes chromatin accessibility, we next examined ATAC-Seq signal at the newly acquired SOX2 sites. Regions gaining SOX2 binding in the absence of CHD4 also showed increased accessibility (Fig. 3E). Indeed, following CHD4 depletion we observed almost 30,000 differentially opening regions that did not correlate with the binding of CHD4 prior to its degradation (Fig. S3A; Table S6), nor do peaks where CHD4 was present show a decrease in H3 occupancy upon its degradation (Fig. S3B). The stronger correlation between accessibility and SOX2 binding— compared with CHD4 (Fig. S3C)—suggested that SOX2 binding likely precedes and facilitates chromatin opening rather than resulting from it. Moreover, footprint enrichment analyses of all regions opening in response to CHD4 degradation, suggest those that are not bound by SOX2 are landing sites of other pluripotency related pioneer factors (Fig. S3D). Taken together these observations suggest CHD4 retains excess TF at sites of active transcription and in its absence, these TFs are free to colonize their higher affinity sites elsewhere in the genome.

To explore the transcriptional potential of these newly accessible sites, we analyzed POLII occupancy at the same regions (Fig. 3E). We observed that POLII was recruited to the new SOX2-bound and ATAC-opening sites, indicating their engagement with the transcriptional machinery. Gene-ontology analysis of the closest genes to peaks that gain SOX2 and POLII binding upon CHD4 degradation shows enrichment of terms related to neural differentiation (Fig. S3E). However, while SOX2 binding and chromatin accessibility remained largely unaffected by ERK activation, POLII occupancy at these regions decreased upon ERK stimulation (Fig. 3E). This observation suggests that the altered nuclear and chromatin landscape in the absence of CHD4 result in ectopic transcription that then is a substrate for ERK mediated repression. Taken together, these observations suggest CHD4 restrains SOX2 binding to cell type-specific sites impeding transcriptional from non-canonical recruitment of POLII by pioneer factors (Fig. 3F).

### CHD4 molds the physiological ERK transcriptional response

Given that CHD4 loss altered SOX2 binding and led to an ectopic transcriptional response to ERK signaling, we next investigated whether it also impacts canonical ERK-induced transcription. To gauge the significance of mutating NuRD components on signaling response, we performed RNA-Seq following 8 hours of cell intrinsic ERK induction. In addition to CHD4 we also mutated other non-canonical components of the NuRD complex associated with MED24 (Fig. S4A). While mutation of SALL1 and SALL4 on their own had a negligible phenotype, signaling failures began to be observed in the compound mutant, and the combination of the SALL1/4 double mutant and CHD4 degradation, substantially hampered the canonical ERK transcriptional response (Fig. S4B). As the 8h phenotype could be masked by secondary effects of the loss of the NuRD complex and its co-factors, we focused on immediate early transcriptional responses to signaling and assessed nascent transcription using TT_chem_-seq^34^ for 45 min and 2 hours following signaling activation (Fig. 4A). We chose 45 min as the first time point as the cRAF module is activated in 30 minutes after 4OHT application and thus this represents 15 minutes post ERK entry into the nucleus^17,35^. Based on principle component analysis (PCA) (Fig. 4B), the largest effect on nascent transcription was ERK signaling, with NuRD depletion resulting in a comparatively modest alteration to the ERK response. Loss of SALL1 had little influence on transcriptional response, while loss of SALL4 and the double mutant resulted in moderate changes. Loss of CHD4 produced the most significant alterations. Despite the good correlation between genes expressed in the 8h RNA-Seq and 2h TT_chem_-Seq data (Fig. S4C, R=0.88, p<2.2e-16), shorter time point nascent transcription suggested that predominant immediate early impact on ERK came from CHD4 itself. In line with this, one of the biggest effects on ERK-dependent transcription was found in the absence of CHD4 alone, where CHD4 had a more significant impact on inductive transcription in response to ERK (Fig. 4C). Individual genes *Dusp5* and *Klf4* followed similar dynamics as the groups of induced and repressed genes (Fig. 4D). To better understand how the NuRD complex affects ERK-driven gene transcription, we analyzed both ERK dependent changes in accessibility and the average nascent transcription over gene bodies of known ERK-regulated genes (Fig. S4D; Table S7). Unlike the reported role of the NuRD complex to act as a chromatin remodeling repressor we observed that NuRD was not required for ERK dependent changes in accessibility, but was required for efficient induction of nascent RNA, but not repression.

**Figure 4.**
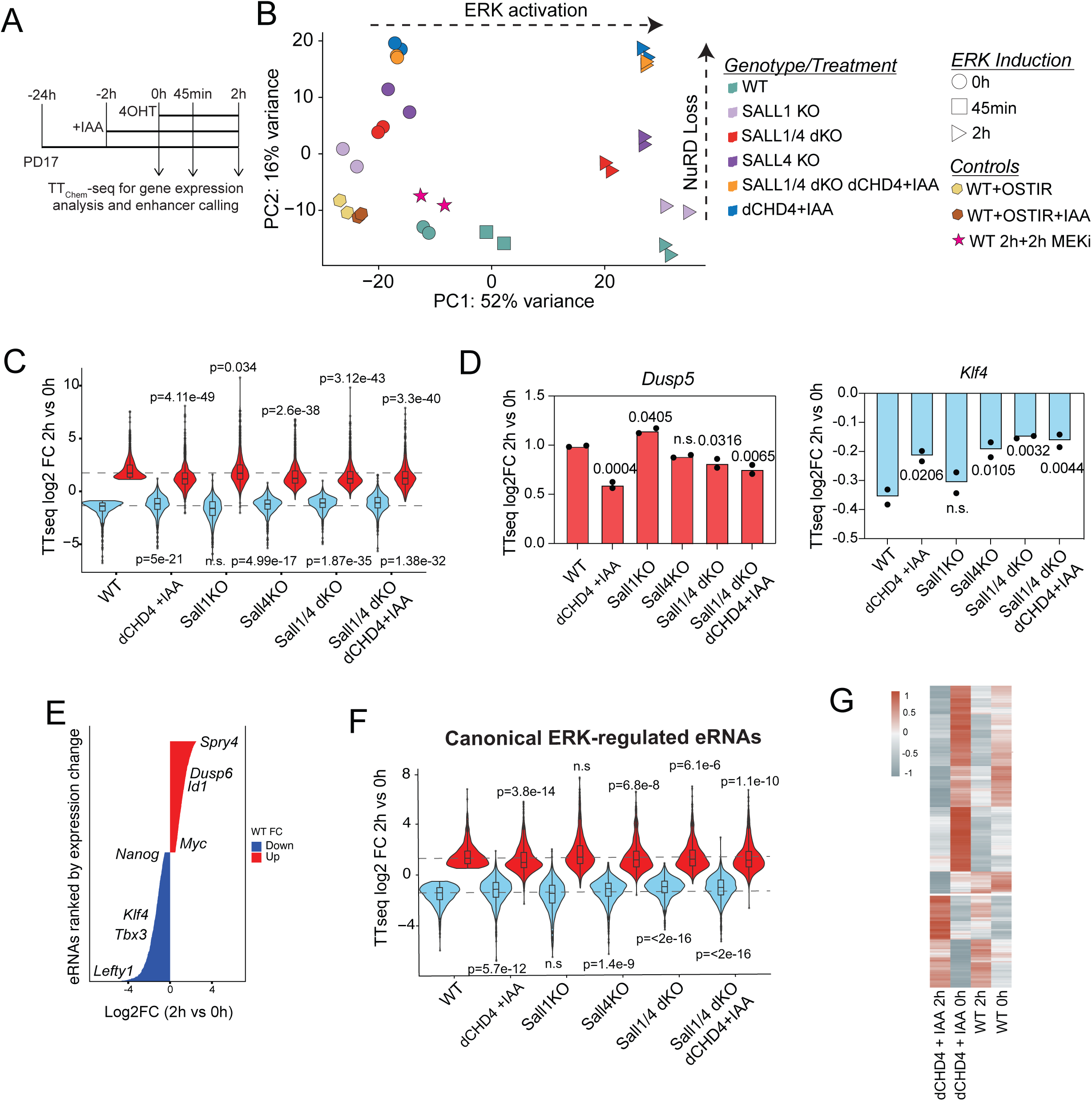
The nucleosome remodeler subunit of NuRD, CHD4 is crucial for ERK-regulated transcription and enhancer activity. **(A)** Schematic showing the experimental setup to access nascent transcription with TT_chem_-Seq upon ERK activation. **(B)** Principal component analysis (PCA) of the nascent transcription data showing how the conditions separate based on their transcriptome upon ERK activation and loss of NuRD, SALL1 or SALL4. PC1 explains 52% of the total variance corresponding to the changes from ERK activation, while PC2 explains 15% of the total variance and separates the different genomic mutations or protein-degradation phenotypes. **(C)** Violin plots representing the distribution of nascent transcription log2 fold-changes (2h of ERK activation vs PD17) of canonical ERK regulated genes (calculated in wild-type (WT) i.e. CHD4-AID not treated with iAA) at the different conditions and genotypes. ERK-repressed genes (blue) n=772, ERK-induced genes (red) n=703. **(D)** Average log2 fold expression between ERK and PD17 treated cells at the different mutant and degron cell lines for two ERK-regulated genes; *Dusp5* and *Klf4*, an induced and repressed gene, respectively. **(E)** Newly characterized ERK-regulated eRNAs ranked by fold change upon 2h of ERK induction, some annotated with closest known ERK-regulated gene. **(F)** Violin plots representing the distribution of nascent transcription log2 fold-changes (2h of ERK vs PD17) of ERK-canonical eRNAs (calculated in WT) at the different conditions and genotypes. ERK-repressed eRNAs (blue) n=678, ERK-induced genes (red) n=662. (**G**) Heatmap of newly ERK-responsive genes in CHD4-depleted cells, identified by differential nascent expression (TT_chem_-Seq) unique to ERK induction. The heatmap displays log₂ fold-change in expression across conditions.

We next wished to understand how the NuRD complex influences enhancer regulation by ERK. To do so, we took advantage of TT_chem_-seq’s ability to capture enhancer RNAs (eRNAs) and augmented our existing set of ERK-dependent enhancers^18^. As described in more detail in the methods, we used GenoSTAN^36^ to define Transcriptional Units (TUs) (Fig. S5A and B) within our TT_chem_-seq data. We used the *Nanog* locus, to exemplify that the genome segmentation correctly identifies both its proximal and distal enhancer (Fig. S5C). To check the quality of our enhancer list, we determined differentially expressed eRNAs between 0h and 2h of ERK induction, annotated them to the closest gene in either direction or ranked them by log-fold change (LFC) (Fig. 4E). We identified many known ERK-activated and repressed genes that mapped to this redefined enhancer list and the correlation between eRNA expression and closest gene expression was positive (R=0.22) at 45 minutes and increased at 2h (R=0.41) of ERK activity (Fig. S5D). Like the analyses for gene transcription, we used this newly identified canonical ERK-regulated eRNAs and calculated the fold change of activity upon ERK for each of the conditions (Fig. 4F). The regulation of eRNA followed the same trend as the gene bodies, in that loss of CHD4 with or without loss of SALL1/4 showed the biggest impairment relative to wild-type (WT). Thus, the rapid degradation of CHD4, the chromatin remodeler component of the NuRD complex is sufficient to affect the ERK canonical transcription response, by affecting both gene and enhancer transcription. Similar to the gene bodies, ERK dependent changes in enhancer accessibility did not dependent on CHD4 (Fig. S4E).

The loss of CHD4 resulted in ectopic binding of SOX2 and POLII recruitment to regulatory regions of neural genes (Fig. S3E). Using the nascent transcriptome, we confirmed that the genes contributing to neural GO terms have a significant increase in expression in the same conditions (Fig. S5E), indicating that loss of CHD4 leads to SOX2 occupying regulatory regions of neural genes and their ectopic expression. Moreover, global analysis of novel ERK-induced transcriptional changes occurring in the absence of CHD4 identified new ERK targets in dCHD4iAA; genes only activated or repressed in the absence of CHD4 (false discovery rate (FDR) > 0.05 and LFC >1) (Fig. 4G). This revealed a large group of genes that were *de novo* repressed, because the level of basal transcription of these genes was reduced in the absence of CHD4, resulting in a low basal expression state that was restored in the presence of ERK signaling.

### Reduced recruitment of pioneer factors through tethering impedes POLII release and eRNA production

As we observed thousands of new SOX2 binding events following CHD4 depletion, we considered that these new events come at the expense of low affinity tethering events that could contribute to ERK responses. Consistent with this interpretation and as shown in Figure 5A, the loss of CHD4 resulted in reduced SOX2 binding at the promoters of impaired ERK-induced genes. Similarly, at ERK-induced eRNAs, where CHD4 depletion resulted in damped enhancer regulation, we observed a reduction in SOX2 recruitment (Fig. 5B). Reflecting our global analysis of ERK mediated SOX2 binding, these SOX2 enhancer regions are enriched for AP1 DNA-binding motifs (Fig. S6A). This suggests that the NuRD complex facilitated the recruitment of SOX2 to low affinity sites in the vicinity of high levels of transcription. This seems to be specific for pioneer factors like SOX2, as SALL4 recruitment is not affected by CHD4 depletion (Fig. S6B). However, like SOX2, SALL1 and SALL4 also exhibit thousands of new binding events upon loss of CHD4, presumably as a result of cooperative binding with other pluripotent pioneer TFs (Fig. S6C).

**Figure 5.**
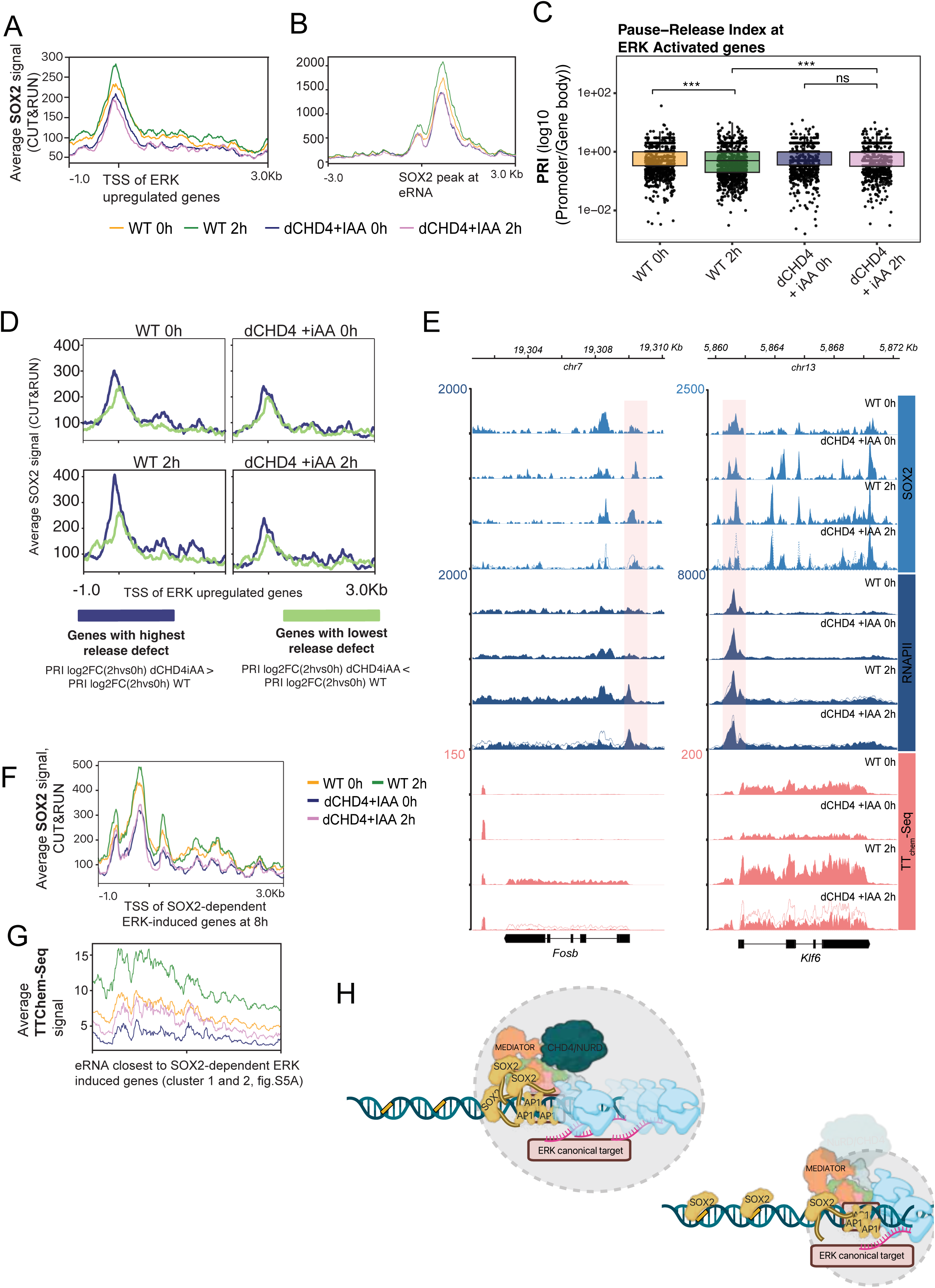
CHD4 sequesters SOX2 at ERK-induced promoters and enhancers to ensure efficient transcription. **(A)** SOX2 occupancy profiles at transcription start sites (TSSs) of ERK-induced genes in control (dCHD4) and CHD4-depleted (dCHD4+iAA) cells before (0h) and after (2h) ERK activation. Lines represent the normalized mean CUT&RUN SOX2 signal around -1Kb and +3Kb of the TSS. **(B)** SOX2 occupancy profiles at ERK induced eRNAs before (0h) and after (2h) ERK activation. Lines represent the normalized mean CUT&RUN SOX2 signal around - 3Kb and +3Kb of the eRNA**. (C)** Boxplot showing the distribution of pause-release index (PRI) values of ERK-induced genes across conditions. Each point represents an individual gene; boxes indicate the median and interquartile range. Statistical significance was assessed using two-sided Wilcoxon rank-sum tests, and p-values were Benjamini-Hochberg adjusted for multiple comparisons. Asterisks denote significance levels: *** p < 0.001, and ns indicates not significant. **(D)** SOX2 CUT&RUN signal stratified by release defect severity. Average SOX2 CUT&RUN signal at ERK-induced gene promoters for the top (left; highest release defect) and bottom (right; lowest release defect) gene groups defined in Fig.S7D. **(E)** Genome browser snapshots illustrating representative examples of genes with higher release defect (e.g., *Fosb*, *Klf6*) with altered SOX2, POLII, and nascent transcription (TT_chem_-seq) signals across conditions (control (dCHD4), iAA-treated (dCHD4iAA), before (0h) and after (2h) ERK). Shaded areas mark regions showing decreased SOX2 binding at the promoter of these genes. In dashed line is represented the tracked of the corresponding control. **(F)** SOX2 occupancy profiles at transcription start sites (TSSs) of SOX2-dependent ERK-induced genes in control (dCHD4) and CHD4-depleted (dCHD4+iAA) cells before (0h) and after (2h) ERK activation. The genes were identified by analysis of prime-seq data of SOX2-depleted cells induced with ERK for 8h (cluster 2 from Fig.S8A). Lines represent the normalized mean CUT&RUN SOX2 signal around -1Kb and +3Kb of the TSS. **(G)** Average TT_Chem_-Seq signal over the eRNAs closest to SOX2-dependent ERK induced genes (clusters 2 and 3 from S8A) (**H**) Schematic for proposed model of action of the NuRD complex of restraining SOX2 binding and promoting robust transcription at signaling responsive regulatory regions.

Given that SOX2 can recruit POLII to sites where it engages with DNA by binding the SOX motif (Fig. 3C and E), we investigated whether the decrease in the tethering-mediated recruitment to signaling responsive sites resulted in decreased POLII. Nascent transcription dynamics over gene bodies indicated that ERK-induced genes have lower transcriptional output when CHD4 is degraded (Fig. S7A, left panel, purple line *vs* green line). We quantified the recruitment of POLII to the promoters of these ERK induced genes by performing ChIP-Seq for total POLII (Fig. S7B). While in the WT there was a significant fold increase in recruitment at 2h of ERK, the levels of POLII recruitment remain high in the absence of CHD4. Moreover, signaling had little effect on these levels, although the overall recruitment to these promoters was the same as that observed with ERK activation in WT. As POLII recruitment was unaffected, despite changes in transcription, we assessed the release of the polymerase into the gene body. To address this, we calculated a Pause-release Index (PRI) per gene based on nascent transcription, that represents the ratio of promoter reads (from -50bp to +1500bp from TSS) to a fixed window of the gene body (from +500bp to +2500bp from the gene body) (Fig. S7C). When comparing the distribution of the PRI values of ERK induced genes in WT and dCHD4 + iAA we observed that while POLII was efficiently released in the WT (0h<2h) this was not the case when CHD4 was absent (0h=2h) (Fig. 5C).

Since CHD4 is an ATP-dependent chromatin remodeler we checked whether the failure to release POLII from the promoter was due to a defect in clearing the +1 nucleosome. However, we found no difference in histone 3 (H3) depletion from this position from ERK-induced genes in response to signaling (Fig. S7D). Together these findings indicate that transcriptional impairment in response to signaling in the absence of CHD4 arose not from reduced POLII recruitment, but from inefficient release into productive elongation, independent of +1 nucleosome clearance.

We next asked whether SOX2 tethering at ERK stimulated active sites is required for efficient POLII release into the gene body and for the transcriptional activation of ERK-induced genes. Initially we classified canonical ERK-induced genes according to the extent to which their promoter-proximal index (PRI) was affected, by comparing the PRI fold change upon ERK induction between WT and dCHD4iAA conditions. Genes showing the strongest release defects displayed higher PRI log₂ fold changes (2h vs 0h) in dCHD4iAA compared to WT cells (Fig. S7E; Table S8A). We then quantified average SOX2 binding at the promoters of genes with the highest and lowest release defects (top and bottom 25% of ranked genes, respectively) (Fig. 5D). Genes with the most pronounced pause–release defect also exhibited the largest reduction in SOX2 promoter binding (Fig. 5D, E).

To directly assess the role of SOX2 tethering ERK-driven transcription, we generated an ERK-inducible cell line in which the endogenous *Sox2* was fused to a FKBP12 (dSox2-cRAF) and selectively degraded SOX2 for 1h prior to inducing the pathway for 8h. Transcriptome analyses revealed that approximately 70% of ERK-induced genes depended on SOX2 (Fig. S8A, cluster 2 (109 genes) and cluster 3 (151 genes); Table S8B). The remaining third of ERK-activated genes, already displayed altered transcript levels before ERK stimulation compared to controls, suggesting secondary effects of SOX2 depletion over the 9-hour period (Fig. S8A, cluster 1 (113 genes). Consistent with what occurs at the promoters with impaired POLII release, in the absence of CHD4, SOX2 recruitment was reduced at promoters of the SOX2-dependent ERK-induced genes Collectively, these findings indicate that tethering-mediated recruitment of SOX2 to promoters and enhancers of ERK-activated genes is essential for their productive transcription. To test for the role of tethering of other pioneer factors in promoting ERK-induced transcription, we employed a double degron for the factors NR5A2 and ESRRB. Like our observations with SOX2, there is a robust reduction of immediate early induction (*Egr1*, *Dusp6* and *Spry4*) by ERK in the absence of these two pioneer factors (Fig. S8B). Based on these observations we conclude that NuRD facilitates signal-dependent transcription by capturing and retaining TFs at cell type–specific target loci, ensuring robust POLII release, eRNA production and a proper transcriptional response to signaling, while protecting the genome from spurious pioneer activity (Fig.5H).

## Discussion

While the canonical view of TF activity is that they are recruited to specific DNA sites and this mediates transcription, here we identify tethering-mediated recruitment of these factors to sites transcription to insure a robust signaling response. We found that ERK mediated phosphorylation of cofactors drives the association of the NuRD complex with the Mediator, simultaneously recruiting a pool of pioneer transcription factors to low-affinity, motif-independent sites of signaling mediated active transcription. At these sites, the concentration of these factors supports productive POLII release and eRNA production. By concentrating these factors at signal-responsive hubs it simultaneously withdraws them from the vast excess of consensus motifs distributed across the genome. In this way NuRD provides an answer to a long-standing paradox: pioneer factors are biochemically capable of engaging chromatin almost anywhere, so why don’t they? Here we suggest that this is because the majority of free TFs are sequestered at active sites of transcription, reducing their concentration elsewhere and also ensuring high local concentrations of TF regulatory domains at sites where signaling drives transcription.

Although the NuRD complex has generally been viewed as a corepressor^25^, we found it is required for optimal signaling response, and that it does so independently of its activity in chromatin remodeling. Despite the diverse enzymatic and chromatin-recognition activities of its subunits^28,29^ ERK-dependent changes in chromatin accessibility persisted even in the absence of CHD4 and disruption of the NuRD complex. However, despite normal signaling dependent chromatin remodeling, ERK mediated transcription is attenuated. This occurs at both the enhancer and promoter level, and at the latter NuRD facilitates ERK mediated pause release. Whereas a role for NuRD in pause release has been previously identified^37^, it was associated with NuRD mediated nucleosome conformation and stability, and we found NuRD depletion produced no reduction in chromatin clearing, suggesting that its role in pause release was unrelated to its capacity to modify chromatin. Similarly, in enhancers, we also observe CHD4 dependent attenuation of transcription, although it is difficult to determine whether this reflects a failure of POLII in eRNA elongation. Although NuRD has been implicated in transcriptional activity, pluripotency exit and lineage commitment, these functions have all been attributed to its activity in chromatin remodeling^38–44^ . Despite having its own remodeling activity, it is also possible that these observations could be mediated via pioneer factor recruitment and their capacity to recruit additional chromatin remodelers in the SWI/SNF family^45,46^ . Moreover, the activity of NuRD in chromatin remodeling has been associated with restricting TF access to DNA^47,48^. However, this would imply that NuRD is bound at these sites keeping them closed, and we observed no correlation between CHD4 binding and regions that open in response to depletion. Instead, we identify DNA binding motifs for pluripotency related pioneer factors in these new accessible regions, *de novo* occupancy by SOX2 complete with POLII recruitment. Thus, the enhanced accessibility accompanying CHD4 loss is likely a consequence of the binding of pioneer factors like SOX2 rather than a direct effect of NuRD.

While SOX2 is the most prominent pioneer colocalizing with CHD4 binding, we also observe sites for canonical pluripotency factors such as OCT4 and KLF4, that could contribute to chromatin opening. Of the pluripotency pioneer factors, SOX2 degradation has the most profound independent impact on chromatin accessibility in mESCs^49^, suggesting it likely plays a primary role in deconstructing the ESC regulon following CHD4 depletion. SOX2 is a pioneer factor expressed in a range of cell types including neural progenitors, anterior foregut, and primordial germ cells^50^. Its capacity to activate neural targets in the CHD4 degron, suggest that loss of its sequestration drives pioneer factor recognition of lineage inappropriate targets. Moreover, in retinal progenitor cells (RPCs) where SOX2 also has a dramatic influence on chromatin accessibility^51^, CHD4 depletion had a similar impact^52^. Thus, as in our observations in ESCs, CHD4 could act in other cell types to restrain SOX2 and other pioneer factors, thereby supporting different cell type–specific enhancer configurations while sustaining a pool of TFs that support plasticity in differentiation.

The failure of ERK mediated induction at promoters was not accompanied by a loss of accessibility or polymerase recruitment, but rather a failure in promoter escape. This defect correlated with reduced SOX2 binding at inducible promoters, suggesting that SOX2 availability at these sites is required to shift paused POLII into productive elongation. Pause-release is known to depend on recruitment and activation of the positive transcription elongation factor (P-TEFb), which can be recruited by and interact with cell-type specific TFs, such as SOX2^53,54^. Here, we propose that the recruitment of SOX2 or other cell type–specific TFs to active transcriptional regions is important for achieving an efficient transcriptional response to signaling by mediating promoter escape and eRNA production. This non-motif-dependent recruitment of SOX2 could be modulated by the presence of intrinsically disordered regions (IDRs) within SOX2 and other Mediator- and NuRD-associated proteins^2,55,56^. These IDRs enable weak, multivalent protein-protein interactions that can retain TFs at high concentration at transcriptional hubs^57^. SOX2 has been shown to form co-condensates with DNA or p300 that enhance transcriptional bursting^58,59^. Furthermore, SOX2 abundance modulates its condensate-forming behavior; when present at low concentration, SOX2 preferentially condenses at high-affinity DNA motifs, but at elevated levels it can also nucleate condensates at lower affinity binding sites^58^.

These observations converge on a concentration-based model for why NuRD is required to constrain SOX2 and other pioneer factors. The genome has far more SOX2 recognition sites than are occupied at any one time, suggesting that when all high affinity sites are occupied the concentration of SOX2 would be insufficient to recognize low-affinity sites where transcription is activated. Our data suggests that NuRD is required to sequester SOX2 at low-affinity transcriptionally active sites to ensure robust transcriptional response and reduce its capacity to freely associate with the thousands of sites containing the TF DNA-binding motif in the genome. In this way, Mediator and NuRD capture sufficient levels of free TFs at sites of transcription to ensure efficient complex formation and exceed the concentration threshold for productive enhancer activation, rapid promoter escape and elongation upon signaling. By doing so, NuRD can impede TF mediated reprograming of somatic cells, but sequestering exogenous reprogramming factors and preventing from accessing new sites^60^. Based on this model, IDR-mediated interactions enable SOX2 and other TFs to be recruited into NuRD-and Mediator-rich hubs, both reducing excess TF to restrain its association with inappropriate enhancer regions and to facilitate productive transcription by increasing its concentration at active regulatory regions

Our findings address why pioneer factor, though able to engage chromatin irrespective of its accessibility, bind only a fraction of their potential sites. By sequestering SOX2 and related factors, the NuRD/Mediator complex simultaneously exploits these factors to obtain an optimal transcriptional output and supports a pool of TFs to enable flexibility in differentiation. At the same time, this excess of factor is withheld from the genome at large - a mechanism that reconciles the promiscuous binding potential of these factors with the precision of signal-responsive gene control.

## Acknowledgments

The authors thank Anja Goth, Jan Zylicz, Joost Gribnau and the entire Brickman lab for critical discussions of this manuscript. H. Wollmann, M. Michaut, A. Kalvisa, and the reNEW Genomics Platform for technical expertise, support, and use of instruments. Gelo dela Cruz and the reNEW Flow cytometry platform for training and support. Smaragda Kompocholi and Lea H. Gregersen for technical support, expertise and reagents for TT_chem_-Seq. Anja Groth and her laboratory, particularly Robert Ciaran MacKenzie Frater for support with ChIP-seq, and valuable feedback along the way. William Hamilton for support in generating the TBR cell line. Naz Salehin and Martin Proks for support in generating and analyzing data for prime-seq. The cell line SOX2-FKBP ESCs was a kind gift from Elzo de Wit lab. Members of the Zylicz and Brickman laboratories for fruitful discussions and critical comments. Work in the Brickman lab was funded by the Lundbeck Foundation (R370-2021-617, R198-2015-412, R400-2022-769, and R286-2018-1534), the Independent Research Fund Denmark (DFF-8020-00100B, DFF-0134-00022B, and DFF-2034-00025B), the Danish National Research Foundation (DNRF116), the Novo Nordisk Foundation (NNF210C0070898), and the European Union (ERC, SENCE, AdG 101097979). The Novo Nordisk Foundation Center for Stem Cell Medicine (reNEW) is supported by the Novo Nordisk Foundation grant number NNF21CC0073729. A. Sell is the recipient of a fellowship from the Novo Nordisk Foundation as part of the Copenhagen Bioscience Ph.D. Programme, supported through grant NNF20SA0035584. J.V.O. acknowledges funding from the Novo Nordisk Foundation under grants no. NNF14CC0001 and NNF24SA0098829. This project was also supported by a generous grant from the Danish Agency of Higher Education and Science to establish the PLATO research infrastructure: Danish National Mass Spectrometry Platform for Proteomics and Biomolecular Imaging (grant no. 5229-00012B).

## Author contributions

Conceptualization, R.S.M, A.C.S., K.B.E., J.V.O. and J.M.B.; Methodology, R.S.M, A.C.S., K.B.E., C.M.S., S.R., F.D., J.V.O. and J.M.B.; Investigation, R.S.M, A.C.S., K.B.E., C.M.S., S.R., and F.D.; Formal Analysis, R.S.M, A.C.S., K.B.E., and S.L.L.; Data Curation, R.S.M, A.C.S., K.B.E., and S.L.L.; Visualization, R.S.M, A.C.S., K.B.E., and S.L.L.; Writing – Original Draft, R.S.M, A.C.S., J.V.O. and J.M.B.; Supervision, J.M.B., E.H., and J.V.O.; Funding Acquisition, R.S.M, A.C.S., J.V.O., and J.M.B.

## Methods

### mESC culture

mESCs were routinely cultured in 0.1% gelatine-coated cell culture plates (Corning) in serum/LIF media consisting of Glasgow Minimum Essential Medium (GMEM, G5154), 10% fetal bovine serum (FBS), 1× MEM non-essential amino acids, 2 mM L-glutamine, 1 mM sodium pyruvate (all from Gibco), 100 μM 2-mercaptoethanol (Sigma), and 1000 U/ml LIF (prepared in house). For all standard culture except ERK induction, 1.6 μM Gsk3-inhibitor (Chir99021: Axon Medchem) was added to the medium. The cells were passaged every 2-3 days.

### Generation of ERK-inducible cell line

An ERK inducible mESC line (TBR) was generated by integrating the ERK-inducible construct^17^ into the *tigre* locus, a safe harbor region that allows for stable and high expression of transgenes in mESCs^61,62^. The sequence of the ERK inducible construct was subcloned from the pBXBSSIP_RTTA plasmid (in house, Brickman lab) to obtain a pCAG-*cRaf-Ert2-T2A-rtta-IRES-Neomycin* sequence. The homology arms for the *tigre* locus were added by following a published targeting strategy^62^.. The sequence of the ERK inducible construct was subcloned from the pBXBSSIP_RTTA plasmid (in house, Brickman lab) to obtain a pCAG-*cRaf-Ert2-T2A-rtta-IRES-Neomycin* sequence. The homology arms for the *tigre* locus were added by following a published targeting strategy^62^. The pCAG-*cRaf-Ert2-T2A-rtta* sequence was cloned into the backbone of pEN396 - pCAGGS-Tir1-V5-2A-PuroR TIGRE donor was a fit from Benoit Bruneau (Addgene plasmid # 92142). For targeting, two different sgRNAs were used, which were cloned into the Cas9-containing plasmid pX458 (Addgene plasmid, # 48138). Sequences were ordered as single-stranded DNA, which were annealed at 95°C for 10 min followed by cooling down to 25°C over 2 min: tigre_sgRNA_1 (fw: caccACTGCCATAACACCTAACTT and rev aaacAAGTTAGGTGTTATGGCAGT) and tigre_sgRNA_2 (fw: caccCTGCCATAACACCTAACTTT and rev: aaacAAAGTTAGGTGTTATGGCAG). For targeting, 1 x 10^6^ E14 mESCs (E14Tg2a subclone 129/Ola^17^) were transfected with a mix of 0.5 μg sgRNA plasmids and 2.5 μg of linearized donor plasmid diluted in Opti-MEM™ (Thermo Fisher Scientific,11058021) using Lipofectamine 2000 (Thermo Fisher Scientific, 11668027). Single clones were selected with neomycin over a time of 8 - 10 days, whereby a concentration of 250 ng/ul was used in the first 3 days and changed to 100 ng/ul for the remaining days. Clones were screened by PCR to verify correct targeting by using one primer pair annealing within the homology arms (*fw:* AGGTCCCCCAGTGAACTCACAGTGGCC, *rev:*TGAGTTCAGCACTTTCCGGGCCTGCC) to amplify a short WT band and another primer pair (*fw:*GACGAAGAGCATCAGGGGCTCGCGCCA, *rev:* TGAGTTCAGCACTTTCCGGGCCTGCC) to amplify the region spanning from the neomycin resistance cassette to the 3’ homology arm to validate insertion into the *tigre* locus. The functionality of the TBR line was tested based on a time course experiment whereby cells were treated with tamoxifen and collected for western blot analysis (Fig. S2B). ERK induction was validated by detecting the phosphorylated form of ERK using the Phospho-p44/42 MAPK (Erk1/2) (Thr202/Tyr204) (Cell signaling, 4370). Phosphorylation of ERK was compared relative to the total ERK using p44/42 MAPK (Erk1/2) (L34F12) (Cell Signaling, 4696) and histone H3 (Abcam, 10799) was used as a loading control.

### Generation of degradation-inducible cell lines

#### dCHD4

To generate an auxin-inducible degron line, mESCs carrying the c-RAF kinase construct in the *tigre* locus, were first lipofected with piggybac-OSTIR-puro plasmid for random integration and selected with puromycin (1µg/ml) for 5 days. The two plasmids targeting the CHD4 locus (pFD104 and pFD105, see below) were co-transfected, 3µg each and seeded in clonal density in medium containing Blasticidin (10 μg/ml) and selected for 10 days. Knock-in clones were identified via PCR and Sanger sequencing.

Primers and PCR settings as follows: CHD4 forward GGAGACAATAGTGAGAAGGAATAATG; CHD4 reverse CACACTCACCCACACAGAACC (WT band of 1129bp and knock-in band of 2034bp)

Using Quick-Load Taq 2X Master Mix (NEB, # M0271) and adding to thermocycler at 95°C for 30 sec, 30 cycles at 95°C for 30 sec, at 50°C for 30 sec, and at 68°C for 90 sec) and final extension at 68°C for 5min.

#### dMed24

To generate an auxin-inducible degron line, E14 mESCs were transfected with a gRNA, targeting the last codon of *Med24,* and a donor template containing a 5’-minAID-eGFP-3’. The construct was generated by amplifying a 5’ homologous 3kb fragment, a 2.8Kb 3’ homologous fragment from a bacterial artificial chromosome (BAC) and the miniAID-GFP was amplified from an in-house plasmid (kind gift from William Hamilton). The gRNA was cloned into Px458-mCherry (Addgene, #161974)^63^.The gRNA was cloned into Px458-mCherry (Addgene, #161974)^63^. Cells were transfected with 2.5μg of linearized donor template and 0.5μg of circular gRNA with the use of Lipofectamine 2000 (Thermos Fisher Scientific, Cat: 11668030). After lipofection cells were expanded and single cell clones were selected by flow activated cell sorting of cells with high levels of GFP. Positive clones were confirmed by PCR, copy number variation (CNV) analysis, Sanger sequencing and Western Blot. These cell lines further edited by transfecting with a donor template cRAF-erT2-rTA-Neo and a gRNA targeting the *tigre* locus. The lipofected cells were selected with Neomycin and the single clones were picked and confirmed by PCR and Sanger sequencing. Finally, to enable the degradation of Med24-GFP, the PB-ostir-puro (Addgene #161972) was edited to include the auxin response factor 16 (ARF16) to control for constitutive degradation of mAID-tagged proteins. Cells were co-transfected with the PiggyBac donor plasmid PB-OSTIR-ARF16_IRES-PURO together with a PiggyBac transposase expression vector by lipofection, followed by puromycin selection. Single clones were selected, and integration was confirmed by PCR and Sanger sequencing.

#### dSox2-cRAF

The original SOX2-FKBP dTAG line was received as a gift from the de Wit lab and previously published in ^64^. This cell line was further modified by introducing an inducible cRAF construct by transfecting SOX2-dTAG cells with a donor template cRAF-erT2-rTA-Neo and a gRNA targeting the *tigre* locus. Cells were selected by treatment with Neomycin, clones were picked, screened by PCR and confirmed based on efficient Erk induction. **dSox2-cRAF:** The original SOX2-FKBP dTAG line was received as a gift from the de Wit lab and previously published in ^64^. This cell line was further modified by introducing an inducible cRAF construct by transfecting SOX2-dTAG cells with a donor template cRAF-erT2-rTA-Neo and a gRNA targeting the *tigre* locus. Cells were selected by treatment with Neomycin, clones were picked, screened by PCR and confirmed based on efficient Erk induction.

### Generation of knockouts

sgRNAs were designed using multilayered Vienna Bioactivity CRISPR (VBC) score^65^. sgRNAs were cloned into a Cas9-2A-GFP plasmid (PX458, Addgene, #48138) using the GeCKO cloning protocol^66^. 1×10^6^ cells were transfected with 1µg plamid. 24h later GFP-positive cells were isolated using FACS and seeded in clonal density. Positive clones were identified via PCR, Sanger sequencing and Western Blot.sgRNAs were designed using multilayered Vienna Bioactivity CRISPR (VBC) score^65^. sgRNAs were cloned into a Cas9-2A-GFP plasmid (PX458, Addgene, #48138) using the GeCKO cloning protocol^66^. 1×10^6^ cells were transfected with 1µg plamid. 24h later GFP-positive cells were isolated using FACS and seeded in clonal density. Positive clones were identified via PCR, Sanger sequencing and Western Blot.

Primers and PCR settings as follows: SALL1 forward CACAAGAAACCCAAGTGGCG; SALL1 reverse GGAAGAGTCGTTGGTCAGCA (WT band of 901bp); SALL4 forward CCTTCCACCACACTCTGACC; SALL4 reverse GGTCGGGAGGCATAAAACCA (WT band of 1085 bp). Using Quick-Load Taq 2X Master Mix (NEB, # M0271) and adding to thermocycler at 95°C for 3 min, 35 cycles at 95°C for 60 sec, at 54°C for 30 sec, and at 68°C for 60 sec) and final extension at 68°C for 3min.

### ERK activation

Before ERK-induction, the cells were seeded and cultured for 24h serum/LIF supplemented in 250 nM FGFR-inhibitor PD17 (PD173074, Sigma, P2499). To induce ERK, 4OHT (4-Hydroxytamoxifen, Sigma, H7904) was added directly to the medium to a concentration of 250 nM. To degrade CHD4, freshly dissolved IAA (Sigma, #I5148) was added directly to the medium to a concentration of 100 µM, 2h before inducing ERK. To ensure complete degradation of SOX2-FKBP during ERK Induction experiments, dTAG-13 was added to the culture medium at a final concentration of 1 µM, one hour prior to addition of 4OHT.

### RIME

Rapid immunoprecipitation mass spectrometry of endogenous proteins (RIME) was performed on two independent Med24 knockout (KO) cell lines with doxycycline-inducible Med24-FLAG expression^18^ as described by to Mohammed et al. (2016), using a monoclonal anti-FLAG antibody.

### Mass Spectrometry Data Analysis

RIME pulldown samples were on-bead digested with trypsin (200 ng; Sigma-Aldrich) overnight in 100 mM ABC (ammonium bicarbonate) buffer, pH∼8. The nuclear proteome samples in 4% SDS (sodium dodecyl sulfate), 100 mM Tris pH 8.5, 1 mM DTT (dithiothreitol), and 10 mM CAA (chloroacetamide) were digested overnight using protein aggregation capture (PAC)^67^ on the KingFisher Flex (Thermo Fisher Scientific) platform.The nuclear proteome samples in 4% SDS (sodium dodecyl sulfate), 100 mM Tris pH 8.5, 1 mM DTT (dithiothreitol), and 10 mM CAA (chloroacetamide) were digested overnight using protein aggregation capture (PAC)^67^ on the KingFisher Flex (Thermo Fisher Scientific) platform. Proteolytic digestion was carried out using hydroxyl beads (ReSyn Bioscience) at a protein-to-beads ratio of 1:2, with endoproteinase LysC (FUJIFILM Wako Pure Chemical Corporation) and trypsin (Sigma-Aldrich) in an enzyme-to-protein ratio of 1:500 and 1:250, respectively. Resulting peptide samples were acidified to a final concentration of 1% FA (formic acid), where after 50% pulldown eluate and 750 ng peptide were loaded on Evotips according to the manufactureŕs instructions for subsequent liquid chromatography - mass spectrometry (LC-MS) analysis of RIME and nuclear proteomes, respectively. Peptide mixtures were analyzed on the Evosep One LC system using an in-house packed 15 cm, 150 μm i.d. capillary column with 1.9 μm Reprosil-Pur C18 beads (Dr. Maisch, Ammerbuch, Germany) using the pre-programmed gradient for 60 samples per day (RIME; 21 min gradient) or 30 samples per day (proteome; 44 min gradient). Column temperature was set to 60°C using an integrated column oven (PRSO-V1, Sonation, Biberach, Germany) and interfaced online with the Q-Exactive HF-X mass spectrometer (RIME samples) or an Orbitrap Exploris 480 MS (Thermo Fisher Scientific, Bremen, Germany) (nuclear proteomes) using Xcalibur software. Spray voltage was set to 2 kV, funnel RF level at 40, and heated capillary temperature at 275°C. For RIME samples analyzed in data-dependent acquisition (DDA) mode, full MS resolutions were set to 60,000 at m/z 200 and full MS AGC target was 3e6 with an injection time (IT) of 45 ms. Mass range was set to 350–1400. For every full scan, the 6 most intense ions were fragmented (collision energy 28%). The AGC target value for fragment spectra was set at 1e5. The resolution was set to 30,000 and IT set to 54 ms. All data were acquired using positive polarity. For single-shot nuclear proteome analysis in data-independent acquisition (DIA) mode, full MS resolutions were set to 120,000 at m/z 200 and full MS AGC target was 300% with an injections time (IT) of 45 ms. Mass range was set to 350–1400. For DIA-MS/MS spectra, the AGC target value was set at 1000% using49 DIA windows of 13.7 m/z isolation width scanning from 361 to 1033 m/z with an overlap of 1 Th. MS/MS resolution was set to 15,000 and IT set to 22 ms and normalized collision energy was 27%. All data were acquired in profile mode using positive polarity.

Raw RIME DDA-files were analyzed by MaxQuant software version 1.6.0.17 using the Andromeda search engine^68^.Raw RIME DDA-files were analyzed by MaxQuant software version 1.6.0.17 using the Andromeda search engine^68^. Proteins were identified by searching the HCD-MS/MS peak lists against a target/decoy version of the mouse UniProt protein database ((UP000000589_10090 and UP000000589_10090_additional release 2017_04 with 22,262 and 36,810 entries, respectively). using default settings. Carbamidomethylation of cysteine was specified as fixed modification and protein N-terminal acetylation, and oxidation of methionine were considered as variable modifications. Minimum peptide length was 7 amino acids, “maximum peptide mass” was 7,500 Da and label min. ratio count was 1 and all peptides were used for protein quantification. Peptide spectrum match (PSM), protein, and site FDR was set to 0.01. Everything else was set to default values. Raw nuclear proteome DIA files were searched in Spectronaut version 19 (Biognosys) against the mouse UniProt protein database (UP000000589 release 2022_5, 21,983 entries) supplemented with a contaminant database from MaxQuant (246 entries). Cysteine carbamidomethylation was set as a fixed modification, and N-terminal acetylation and methionine oxidation were set as variable modifications, with a maximum number of variable modifications kept at 5, and a maximum number of missed cleavages kept at 2. Cross-run normalization was allowed. The RIME dataset and single-shot nuclear proteomes were normalized and imputed using Prostar software^69^.The RIME dataset and single-shot nuclear proteomes were normalized and imputed using Prostar software^69^. The datasets were filtered removing contaminants and requiring at least 2 (RIME) or 1 (proteome) quantified value for at least one condition. Global quantile alignment was used for normalization and for missing values imputation, the slsa and det quantile algorithms were used for partially observed values (POV) and missing on entire condition (MEC), respectively. The volcano plots showing differential regulation were generated by plotting the –log_10_ transformed and FDR-adjusted p-values (q-value threshold of 0.05) derived from a two-sided t-test versus log_2_ transformed fold changes. Significance was determined based on a hyperbolic curve threshold of s0=0.1using Perseus^70^.Significance was determined based on a hyperbolic curve threshold of s0=0.1using Perseus^70^.

### CO-IP

dMED24-GFP Cells at 80% confluency in a 10cm dish were washed with phosphate-buffered saline (PBS). Cells were scraped in 300μl of NETN Buffer (100mM NaCl, 1mM EDTA, 20mM Tris-HCl pH8.0, 0.5% NP-40) and collected in a cold tube. This was proceeded by a 30min incubation at 4°C while shaking followed by a centrifugation at 13’000xg for 5min at 4°C. The supernatant was collected and Pierce^TM^ BCA Protein Assay (Thermo Scientific, Cat #23225) was used to quantify the protein, and similar amounts were used for immunoprecipitation. 100μl of Dynabeads^TM^ Protein A (Invitrogen Cat# 10001D) with 10ul of rabbit α-GFP antibody (Abcam, ab290) for 1h at room-temperature and added to the protein lysate for another overnight incubation at 4°C. The beads were washed 3 times with NETN buffer using a magnetic rack and eluted in 50μl of SDS elution buffer (250mM Tris-HCl pH6.8, 4% SDS, 20% Glycerol) by incubating at 50°C for 10min. Beads were centrifuged and collected with magnetic rack and the supernatant transferred to a new tube and DTT (100mM) was added prior to separation and detection of MED24 interacting proteins by Western Blot.

### Western Blot

Cells were lysed on the plate in 4% SDS, 20% glycerol, 120mM Tris pH 7.4 and sonicated and quantified on a Nanodrop2000. 40 µg protein per sample was denatured by heat and 100mM DTT. Samples were loaded and resolved on 4-12% NuPAGE Bis-Tris Mini Protein Gels (Invitrogen) in NuPAGE MES SDS Running Buffer (Invitrogen, #NP0002). They were transferred to nitrocellulose membranes (GE Healthcare) via wet transfer in 25mM Tris, 192mM glycine and 20% methanol. The membranes were blocked for 2h at room temperature in 10% skim milk in 150mM NaCl, 10mM Tris, 0.1% Tween (TBST). Respective primary antibody incubation was performed over night at 4°C in 5% BSA in TBST. They were detected by conjugated secondary antibodies (Alexa Flour, Molecular Probes), 2h incubation at room temperature, and visualized using fluorescence in a Chemidoc MP (BioRad).

### CUT&RUN

CUT&RUN^31^ was performed on adherent cells attached to 24-well cell culture plates as previously described^71^ and using an in house purified pA/G MNase^72^. Libraries were generated with the NEBNext® Ultra™ II DNA Library Prep Kit for Illumina (New England Biolabs, Cat. #E7645) and were pooled to equal molarity and paired-end sequenced on Illumina NextSeq 2000 Sequencer according to the manufacturer’s instructions. Sequencing data was processed using the nf-core/cutandrun pipeline^73^ with the following parameters: genome: mm10, trim_nextseq: 20, normalisation_mode: RPKM, extend_fragments: true, normalisation_binsize: 1, seacr_stringent: stringent, seacr_norm: norm, skip_heatmaps: true, dedup_target_reads: true, save_spikein_aligned: false. For differential binding analysis was done with the R package DiffBind^74^, peakset comparisons were done using BEDTools^75^, coverage heatmaps and plots were generated with deepTools^76^ and tracks were created with pyGenomeTracks^77^. *De novo* motif analysis for CHD4 peaks was performed using Hypergeometric Optimization of Motif Enrichment (HOMER)^78^.^71^ and using an in house purified pA/G MNase^72^. Libraries were generated with the NEBNext® Ultra™ II DNA Library Prep Kit for Illumina (New England Biolabs, Cat. #E7645) and were pooled to equal molarity and paired-end sequenced on Illumina NextSeq 2000 Sequencer according to the manufacturer’s instructions. Sequencing data was processed using the nf-core/cutandrun pipeline^73^ with the following parameters: genome: mm10, trim_nextseq: 20, normalisation_mode: RPKM, extend_fragments: true, normalisation_binsize: 1, seacr_stringent: stringent, seacr_norm: norm, skip_heatmaps: true, dedup_target_reads: true, save_spikein_aligned: false. For differential binding analysis was done with the R package DiffBind^74^, peakset comparisons were done using BEDTools^75^, coverage heatmaps and plots were generated with deepTools^76^ and tracks were created with pyGenomeTracks^77^. *De novo* motif analysis for CHD4 peaks was performed using Hypergeometric Optimization of Motif Enrichment (HOMER)^78^.

### TT_chem_-seq

TT_chem_-seq was performed as previously described^34^. Samples were sequenced single-end on an Illumina NextSeq 2000 Sequencer according to the manufacturer’s instructions. Adapters were trimmed from paired-end reads using Cutadapt v4.5^79^ then RiboDetector ^80^ was used to remove rRNA-related sequences. Remaining reads were aligned to the mouse mm10 genome and counts at gene bodies produced using STAR 2.7.9a^81^. Bam files were filtered using SAMtools to keep only alignments with MAPQ of 7 or greater, and PCR duplicates were removed using UMI-tools^82^. Differential expression analysis was performed using DESeq2^83^.Adapters were trimmed from paired-end reads using Cutadapt v4.5^79^ then RiboDetector ^80^ was used to remove rRNA-related sequences. Remaining reads were aligned to the mouse mm10 genome and counts at gene bodies produced using STAR 2.7.9a^81^. Bam files were filtered using SAMtools to keep only alignments with MAPQ of 7 or greater, and PCR duplicates were removed using UMI-tools^82^. Differential expression analysis was performed using DESeq2^83^.

### Enhancer Calling on TTchem-seq data

TU filter^84^ was first used to find intervals of uninterrupted transcription in our TT-seq data.TU filter^84^ was first used to find intervals of uninterrupted transcription in our TT-seq data. The genome was binned into consecutive 200 bp bins and genome-wide coverage calculated, then a hidden Markov model with Poisson Log-Normal emission distribution was fitted using GenoSTAN^36^ to segment the genome into transcribed and un-transcribed states. Each transcribed unit was classified either as protein-coding or as one of several ncRNA categories. Chromatin state segmentation was subsequently performed using GenoSTAN to learn a hidden Markov model with multivariate Poisson-Log-Normal distributions to define putative enhancer regions based on ATAC-seq and patterns of histone modifications (H3K27ac, H3K4me1, and H3K27me3) and EP300 binding. Chromatin states were manually annotated as either active, repressed, inaccessible, or low signal based on the chromatin segmentation results, then transcribed ncRNAs (excluding downstream-sense RNAs) at 0h and 2h Erk induction that originated from enhancer regions that were active at the same timepoint were classified as eRNAs. BEDTools *multicov* was used to produce stranded read counts at these eRNAs and differential expression analysis was done using DESeq2.

### Bulk RNA-seq

2.5×10^4^ cells/cm^2^ were seeded 24h before inducing ERK for 8h. Cells were lysed and total RNA was isolated with the RNeasy Mini Kit (Qiagen, #74104) and 1 µg of total RNA was used for library preparation using the NEBNext Poly(A) mRNA Magnetic Isolation Module (NEB #E7490) and NEBNext^®^ Ultra^™^ II Directional RNA Library Prep kit (NEB, # E7765). Samples were single-end sequenced on an Illumina NextSeq 2000 Sequencer according to the manufacturer’s instructions. bcl2fastq (v2.19.1) was used for raw read processing, STAR (v2.5.3a) for mapping of reads and generation of the count table^81^. DEseq2^83^ was used for determining differentially expressed genes and to generate PCAs in R. The significance cutoff was set at LFC>1 and padj<0.05.2.5×10^4^ cells/cm^2^ were seeded 24h before inducing ERK for 8h. Cells were lysed and total RNA was isolated with the RNeasy Mini Kit (Qiagen, #74104) and 1 µg of total RNA was used for library preparation using the NEBNext Poly(A) mRNA Magnetic Isolation Module (NEB #E7490) and NEBNext^®^ Ultra^™^ II Directional RNA Library Prep kit (NEB, # E7765). Samples were single-end sequenced on an Illumina NextSeq 2000 Sequencer according to the manufacturer’s instructions. bcl2fastq (v2.19.1) was used for raw read processing, STAR (v2.5.3a) for mapping of reads and generation of the count table^81^. DEseq2^83^ was used for determining differentially expressed genes and to generate PCAs in R. The significance cutoff was set at LFC>1 and padj<0.05.

### ATAC-seq

ATAC-seq was performed as previously described^85^. Libraries were pooled to the same molarity and paired-end sequenced on Illumina NextSeq 2000 Sequencer according to the manufacturer’s instructions. Raw read processing was performed using bcl2fastq and adapter trimming using cutadapt ^79^. Bowtie2^86^ was used for read mapping to mm10. Peaks were called with macs2^87^ (v2.2.7.1). Consensus peaks were defined as peaks present in two out of three replicates using the DiffBind package in R ^74^.The same package was used to determine differentially open peaks with LFC>1 and padj<0.05. Differential transcription factor footprinting was predicted using Transcription factor Occupancy prediction By Investigation of ATAC-Seq Signal (TOBIAS)^88^. For visualizing ATAC-seq fragment density distribution V-plots were generated using the R package VplotR^89,90^.ATAC-seq was performed as previously described^85^. Libraries were pooled to the same molarity and paired-end sequenced on Illumina NextSeq 2000 Sequencer according to the manufacturer’s instructions. Raw read processing was performed using bcl2fastq and adapter trimming using cutadapt ^79^. Bowtie2^86^ was used for read mapping to mm10. Peaks were called with macs2^87^ (v2.2.7.1). Consensus peaks were defined as peaks present in two out of three replicates using the DiffBind package in R ^74^.The same package was used to determine differentially open peaks with LFC>1 and padj<0.05. Differential transcription factor footprinting was predicted using Transcription factor Occupancy prediction By Investigation of ATAC-Seq Signal (TOBIAS)^88^.

### H3 and Total PolII ChIP

H3 and PolII ChIP followed by library preparation, Illumina sequencing and data analysis were performed as previously described^91^ H3 and PolII ChIP followed by library preparation, Illumina sequencing and data analysis were performed as previously described ^91^

### Prime-seq

Prime-Seq was performed as previously described^92^. Libraries were pooled to the same molarity and paired-end sequenced on Illumina NextSeq 2000 Sequencer according to the manufacturer’s instructions. Raw reads were processed using bcl2fastq and counts tables were generated using an in-house nf-core pipeline^93^ with the following parameters: r= v1.1, genome=GRCm39. Differential analysis was carried out using the R package DESeq2^83^. Heatmaps were generated using the ComplexHeatmap R package^94^.Prime-Seq was performed as previously described^92^. Libraries were pooled to the same molarity and paired-end sequenced on Illumina NextSeq 2000 Sequencer according to the manufacturer’s instructions. Raw reads were processed using bcl2fastq and counts tables were generated using an in-house nf-core pipeline^93^ with the following parameters: r= v1.1, genome=GRCm39. Differential analysis was carried out using the R package DESeq2^83^. Heatmaps were generated using the ComplexHeatmap R package^94^.

## Data availability

The mass spectrometry nuclear proteome and RIME proteome data have been deposited with the ProteomeXchange Consortium via the PRIDE partner repository with the dataset identifier PXD073070 (RIME) and PXD073174 (nuclear proteome). The next-generation sequencing data have been deposited in the Gene Expression Omnibus, under the accession numbers GSE314818 (ATAC-Seq), GSE314819 (TT_Chem_-Seq), GSE314821 (RNA-Seq), GSE314822 (CUT&RUN), GSE314823 (ChIP-Seq), GSE318753 (CUT&Tag).

## Supplemental information

**Supplementary Figure 1:**
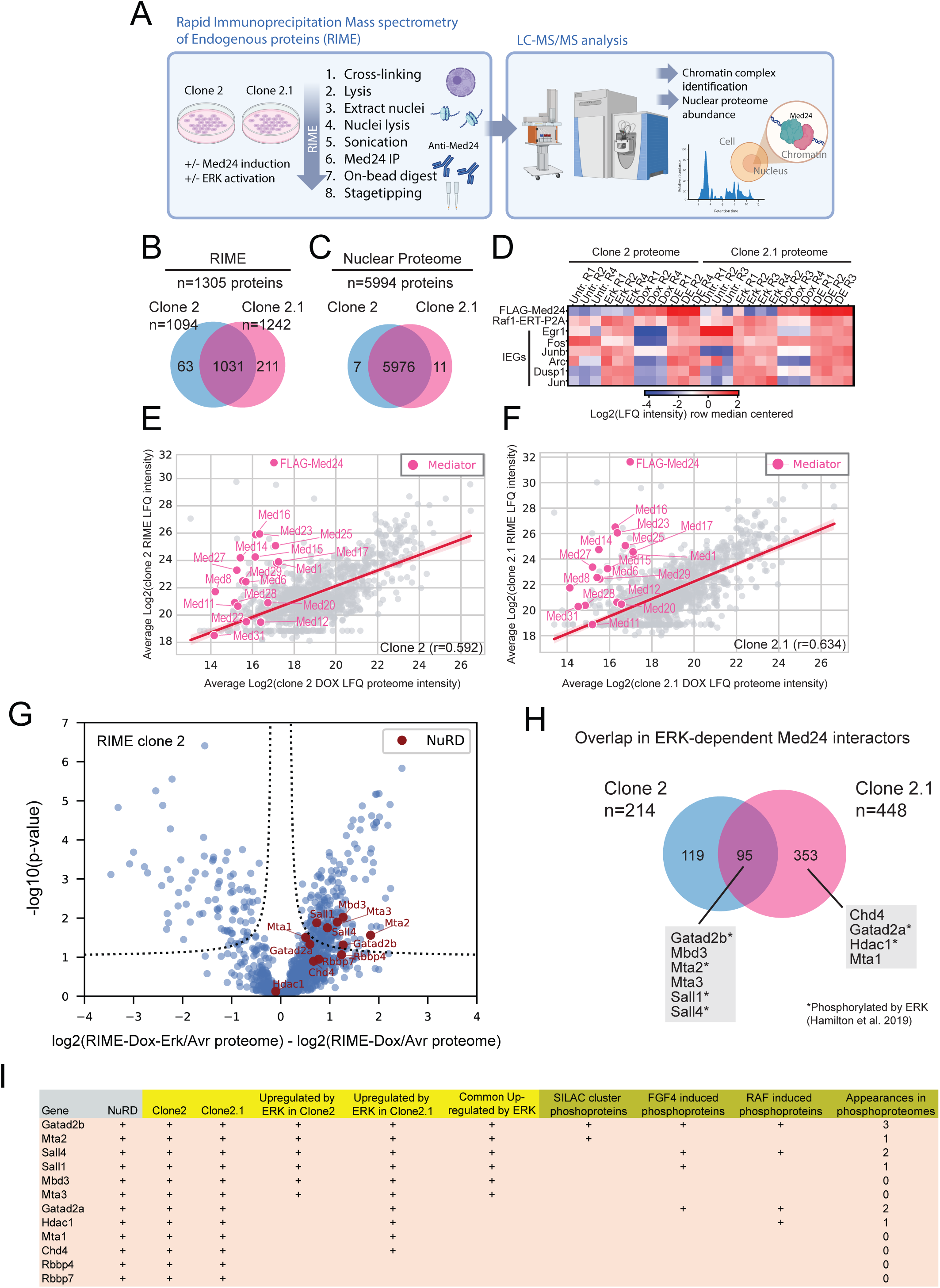
MED24 as a conduit for ERK response associated with the NuRD complex. **(A)**. Schematic representation of the MS-based workflow for rapid immunoprecipitation mass spectrometry of endogenous proteins (RIME) and nuclear proteome analysis. **(B)** and **(C)** Overlap in identified and quantified proteins in the RIME **B** and nuclear proteomes **C** for two analyzed cell clones. **(D)** Heatmap of MED24-induced (dox) and RAF1-induced (4OHT) nuclear abundance and selected immediate early genes across conditions. **(E)** and **(F)** Correlation plot of RIME abundance versus nuclear proteome abundance for Dox-induced conditions in clone 2 **E** and clone 2.1 **F**. **(G)** Volcano plot of RIME data for clone 2 highlighting NuRD complex-related proteins by gene name (red). -log10(p-value) is plotted against log2 fold-change (proteome normalized) for the ERK-induced condition versus control. **(H)** Overlap in ERK-dependent MED24 interactors between the two analyzed clones. **(I)** Table showing NuRD components identified in the RIME for the two clones. Green columns (right) refer to ERK-phosphoproteomes published in Hamilton et al., 2019.

**Supplementary Figure 2:**
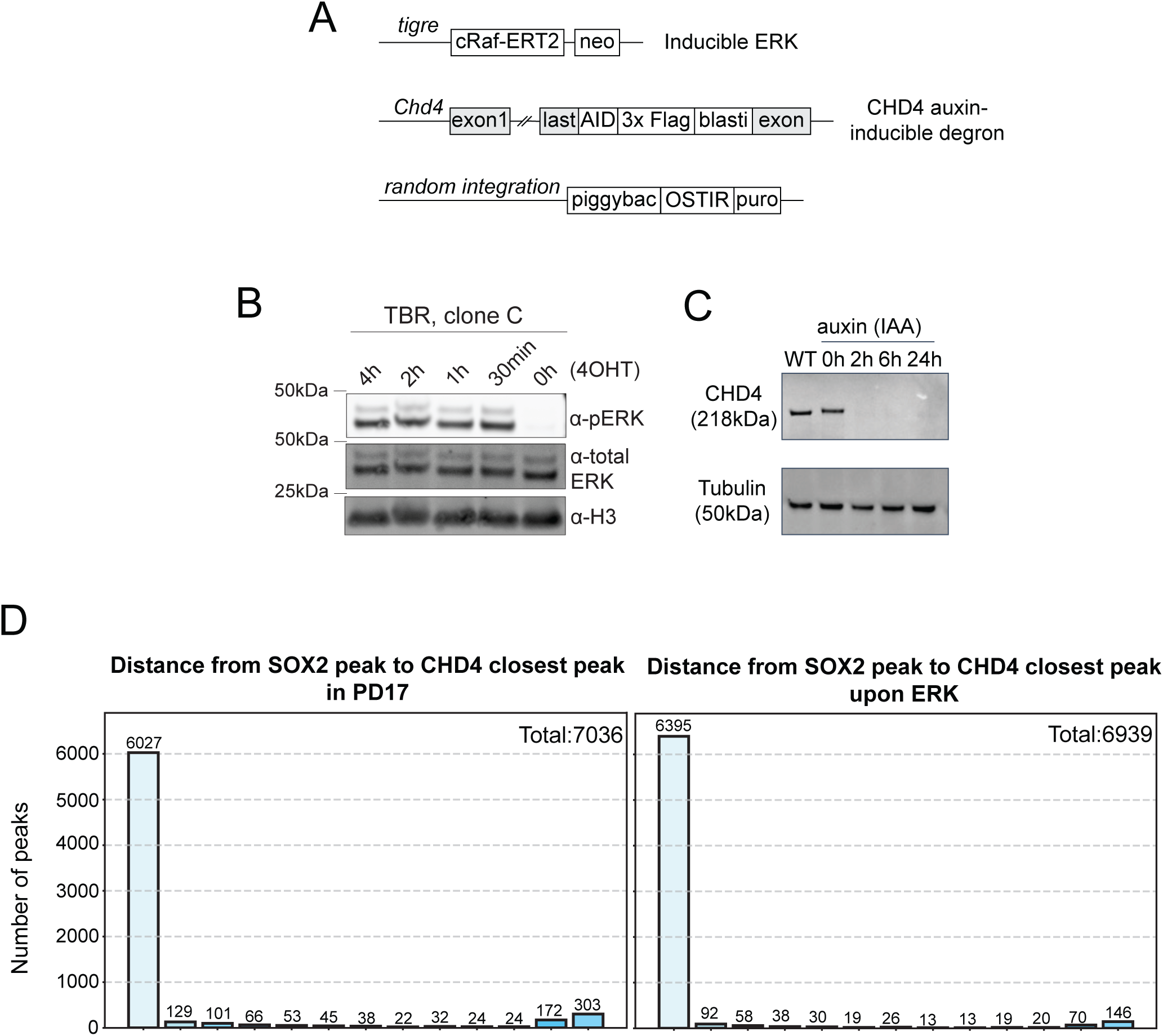
Genetic tools to ablate CHD4 function and explore its relationship to SOX2 binding. **(A)** Schematic representation of genetic modifications used to generate the inducible ERK and auxin inducible CHD4 degron mESC line. **(B)** Western blot screening of ERK-inducible cell line TBR (clone C, used as parental line for dCHD4) by assessing phosphorylated ERK (p-ERK) upon addition of 4OHT. **(C)** Western Blot screening of the auxin-inducible degradation of CHD4 cell line (dCHD4) by assessing loss of the protein upon addition of IAA. **(D)** Distribution of distances between SOX2 and CHD4 peaks closest peaks in the absence of ERK, PD17 (left) and after 2h of ERK activation (right), showing co-localization and increased overlap upon signaling.

**Supplementary Figure 3:**
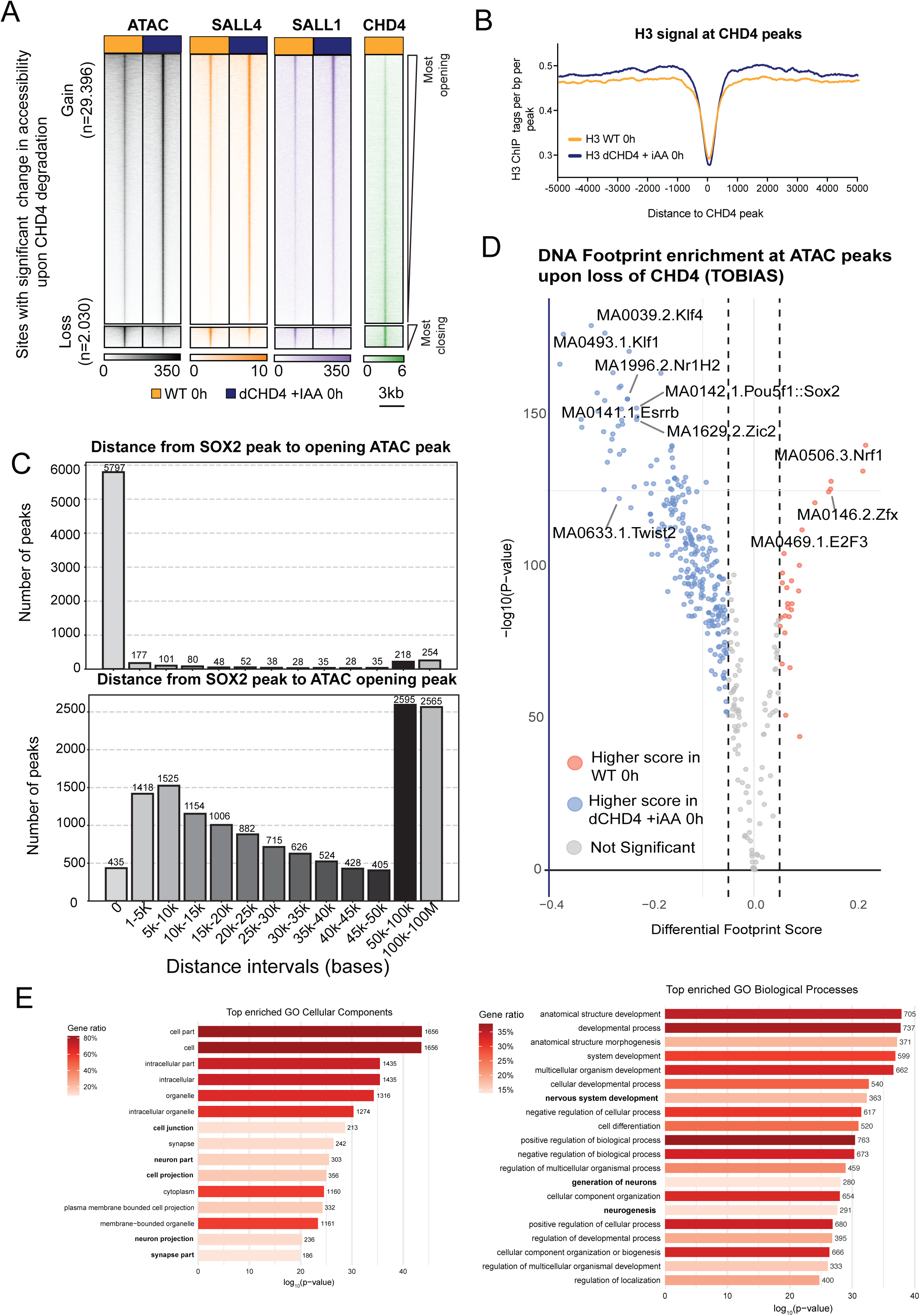
CHD4 depletion results in ectopic SOX2 binding. **(A)** Changes in chromatin accessibility and CUT&RUN signal for SALL1 and SALL4 following CHD4 degradation. The regions represented have significant change in accessibility, measured by ATAC-Seq, upon loss of CHD4 and are ranked by degree of accessibility change. The biggest effect is gain (n = 29,396 sites) of accessibility and smallest effect on loss (n = 2,030). Color scales represent normalized signal intensity, with sites ordered from most opening (top) to most closing (bottom). **(B)** Average H3 ChIP-seq signal centered on CHD4 peaks in control and CHD4-depleted (+IAA) cells without ERK activity (0h or PD17). **(C)** Histogram depcting the distance between SOX2 (top) or CHD4 (bottom) closest peaks to ATAC peaks opening in response to the loss of CHD4. There is a larger overlap of binding of SOX2 (distance =0) to the opening ATAC peaks than with sites where CHD4 binds. (**D**) Volcano plot depicting differential footprint scores and significance (-log10(pvalue) for DNA binding motifs as calculated by TOBIAS. The differential score was calculated on consensus ATAC peaks and accessibility differences in WT vs dCHD4 +iAA prior to ERK activation (0h, PD17). (**E**) Gene Ontology (GO) term enrichment of genes associated with peaks with SOX2 and POLII gain upon NuRD degradation. Top 15 enriched GO (Cellular Components, left and Biological Processes, right) terms for closest genes annotated using HOMER. Terms are ranked by significance (-log_10_ p-value); bar length indicates the enrichment p-value and the fill color indicates the gene ratio (fraction of target genes annotated to each term). The number at the end of each bar is the count of target genes for that term.

**Supplementary Figure 4:**
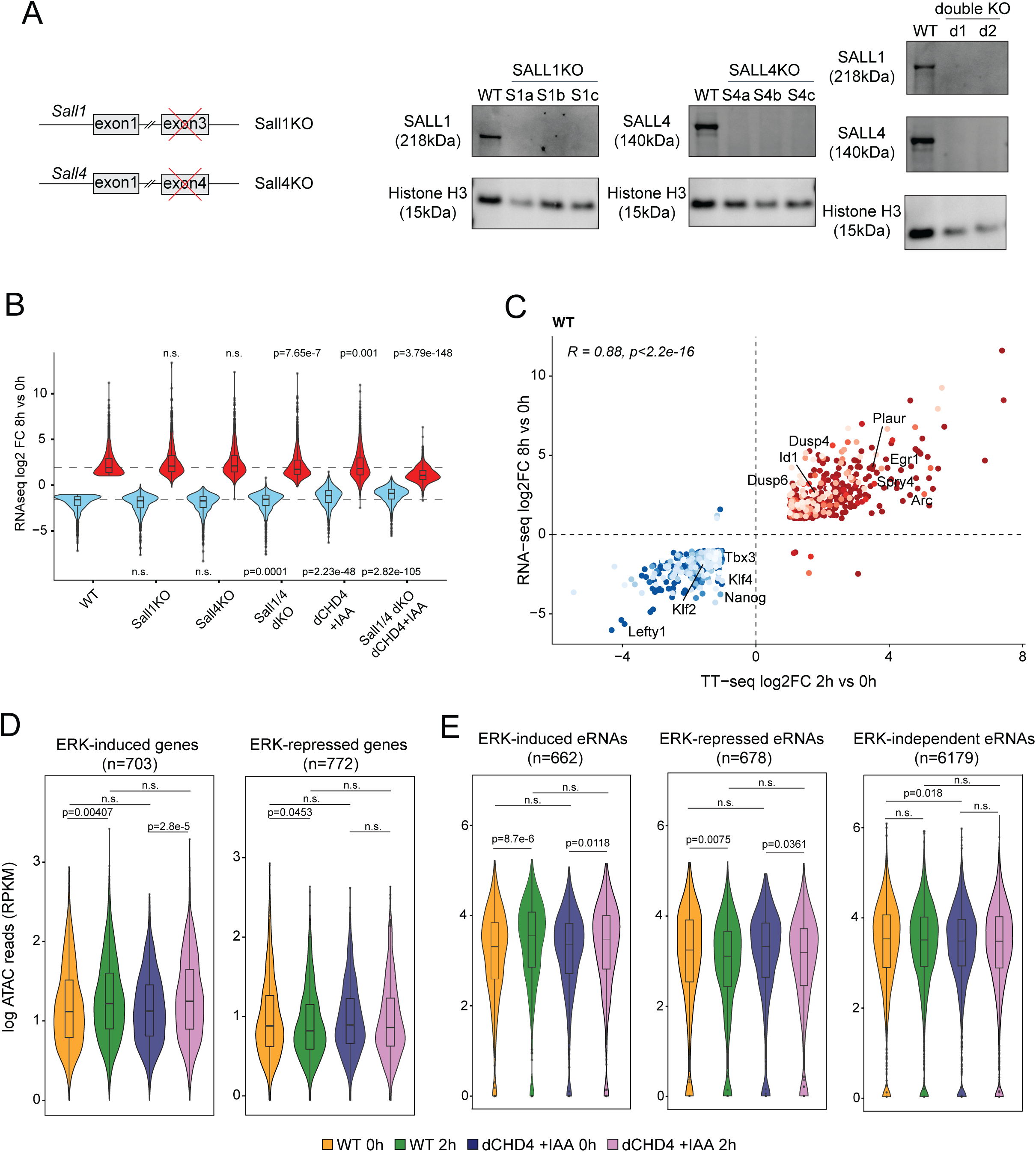
Loss of NuRD affects ERK-driven gene activation independently of co-factors SALL1/4 and accessibility changes. **(A)** Schematic representation of genetic modifications used to generate the SALL1 and/or SALL4 knockouts cell lines in the parental dCHD4, ERK inducible cell line. On the right is the clone screening by Western Blot that confirm the absence of the targeted proteins. (**B**) Violin plots representing the distribution of total transcription log2FC (8h of ERK activation vs PD17 treatment, 0h) of canonical ERK regulated genes in each mutant background shown in **C**. ERK-induced genes are shown in red (n=1221), ERK-repressed genes are shown in blue (n=966). **(C)** Correlation of the log2FC of total RNAseq (8h of ERK activation vs PD17 treatment, 0h) and log2FC of TT-seq (2h of ERK activation vs PD17 treatment, 0h). Each dot represents a gene. ERK induced genes are shown in red; ERK repressed genes are shown in blue. (**D**) Violin plots showing promoter accessibility (log-transformed ATAC-seq reads, RPKM) at ERK-induced (left) and ERK-repressed (right) genes in control and CHD4-depleted cells before (0h) and after (2h) ERK activation. Horizontal lines denote median and interquartile range. p-values correspond to pairwise Wilcoxon rank-sum tests; n.s., not significant. **(E)** Distribution of normalized ATAC-Seq reads (RPKM) at ERK-regulated eRNAs across different conditions.

**Supplementary Figure 5:**
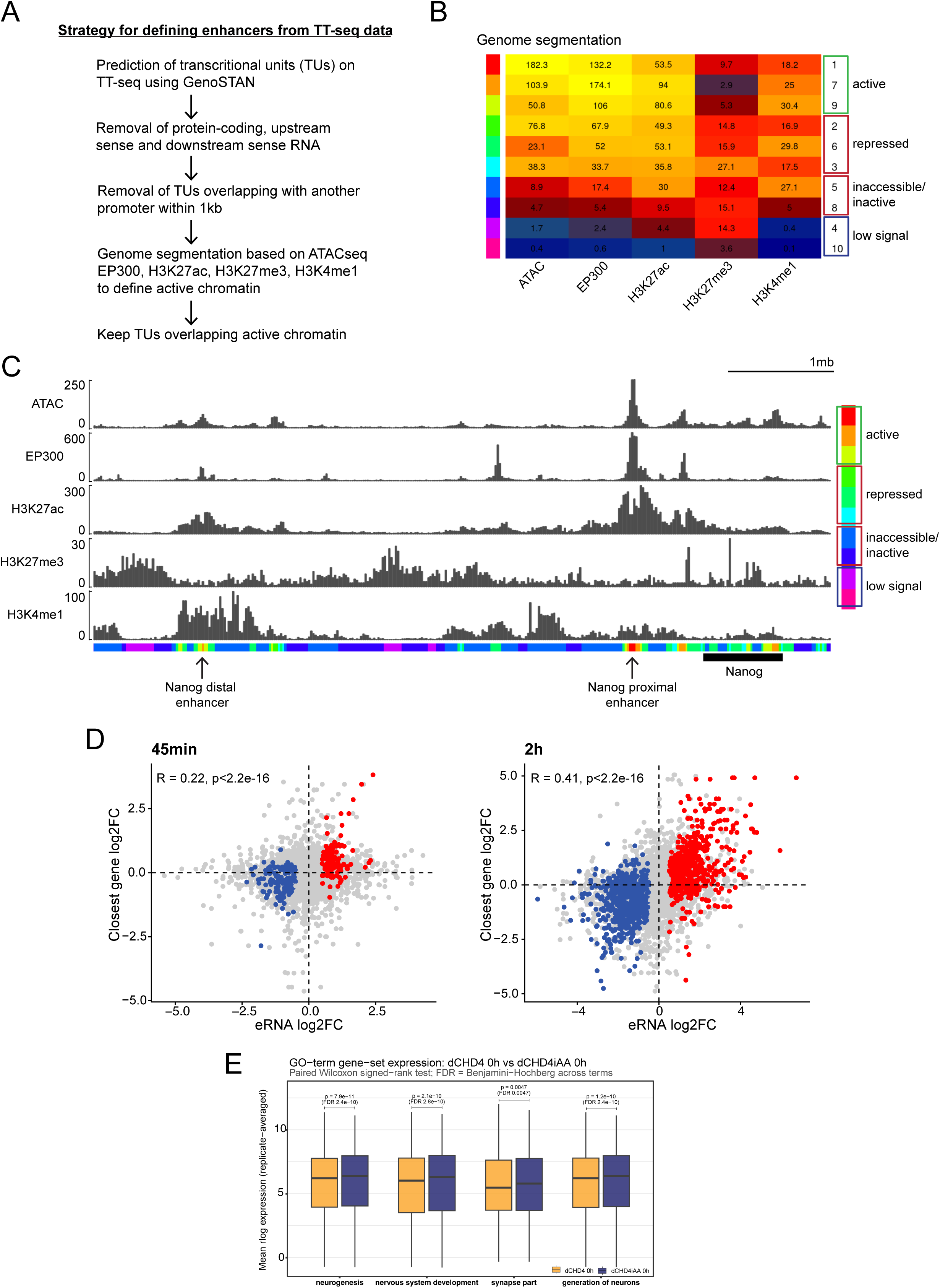
Nascent ERK-dependent transcription. **(A)** Workflow for identification of enhancer RNAs from TT_chem_-seq data. **(B)** Heatmap of genome segmentation scores based on ATAC-seq and EP300, H3K27ac, H3K27me3, H3K4me1 binding. Active chromatin was defined with high scores in ATAC-seq, EP300 and H3K27ac. EP300, H3K27ac, and H3K27me3 data are from Hamilton et al., 2019. **(C)** Genome Browser snapshot showing the region containing the Nanog gene and its proximal and distal enhancer as a representation of correct enhancer identification through genome segmentation. **(D)** Correlation of the log2FC of eRNAs defined by GenoSTAN (45min vs 0h, left; 2h vs 0h, right) and log2FC of genes closest to each eRNA, expression defined by TT_chem_-seq. Each dot represents an eRNA and its closest gene. In red are ERK induced genes; in blue are ERK repressed genes; and in gray are other genes. (**E**) TT_chem_-Seq expression of GO-term sets in dCHD4 versus dCHD4iAA in the absence of ERK (PD17, 0h). Distribution of r-log-normalized TT_chem_-Seq expression, averaged across replicates, for genes annotated for neural-related GO terms selected from the top 15 most enriched as depicted in Figure S3E. Statistical significance was assessed by term by a paired Wilcoxon signed-rank test (paired genes across the two conditions), with p-value shown above each pair and Benjamini-Hochberg FDR-adjusted across the four terms.

**Supplementary Figure 6.**
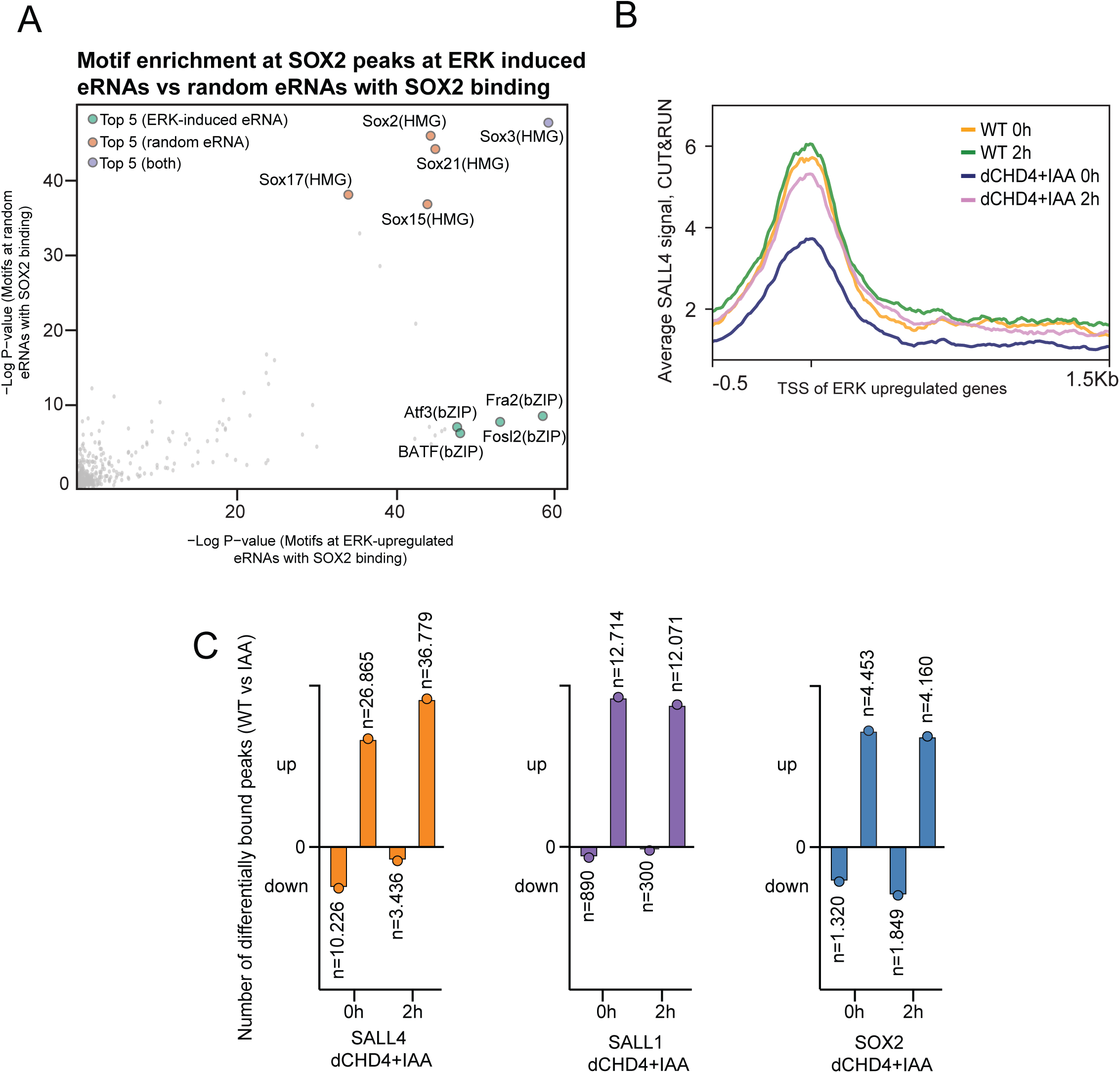
TF localization upon CHD4 degradation. **(A)** Scatter plot comparing the significance of known motifs enrichment at SOX2 peaks overlapping with ERK-induced eRNAs vs random eRNAs. Top 5 known motifs found in ERK induced eRNAs are shown in green, top 5 known motifs found in random eRNAs in orange and common top 5 in blue. Values were calculated using HOMER. **(B)** SALL4 occupancy profiles at TSSs of ERK-induced genes in control (dCHD4) and CHD4-depleted (dCHD4+iAA) cells before (0h) and after (2h) ERK activation. Lines represent the normalized mean CUT&RUN SALL4 signal around -500bp and +1.5Kb of the TSS. **(C)** Bar plots showing absolute numbers of differentially bound peaks (FDR>0.05, log2fc >1) upon loss of CHD4 for SALL4 (orange, left), SALL1 (purple, middle), and SOX2 (blue, right).

**Supplementary Figure 7.**
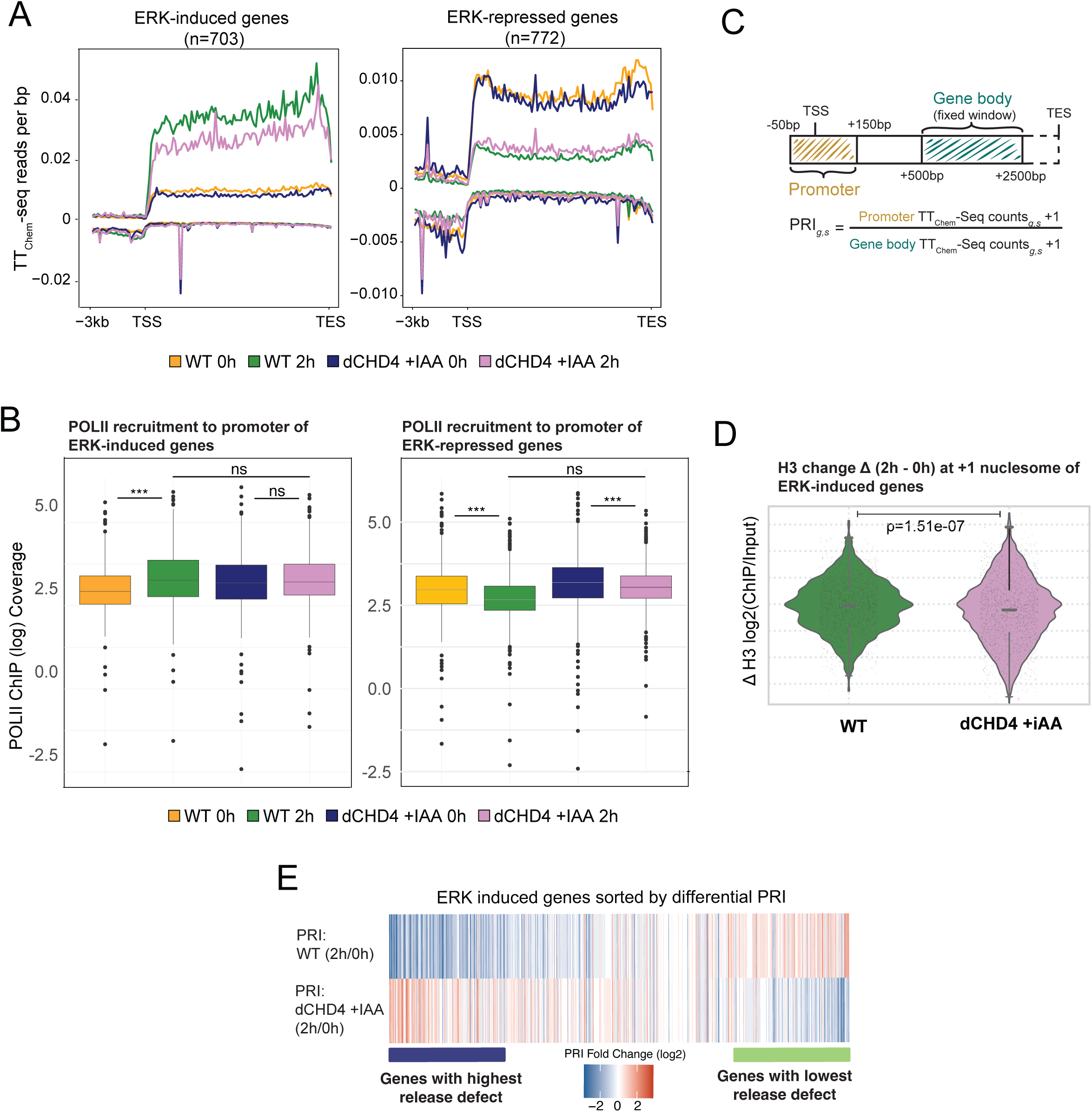
CHD4 is required for efficient POLII release independently of chromatin remodeling. **(A)** Average TTChem-Seq signal over ERK-induced and repressed genes for WT (CHD4-AID not treated with iAA) and dCHD4iAA (CHD4-AID treated with iAA 2h prior to ERK induction) cells before and after 2h of ERK signaling activity. TSS – transcription start site, TES – transcription end site. **(B)** Distribution of POLII ChIP Coverage at promoters of ERK-induced and repressed genes, the same genes as in **A**. **(C)** Schematic of the calculation of the pause-release index (PRI) for gene g in sample s. For each of the ERK induced genes normalized TT_chem_-Seq reads that map to the promoter (-50bp and +150bp from the TSS) and reads that map to a fixed window of the gene body (+500bp to +2500bp from the TSS) were considered for the PRI calculation. **(D)** Violin plots show the distribution of ΔH3 values (log₂[ChIP/Input] difference between 2 h and 0 h) at +1 nucleosome positions (+50bp-+250bp from TSS) at ERK-induced genes for the untreated control (dCHD4) and iAA-induced degradation of CHD4 (dCHD4+iAA). Each point represents the ΔH3 for an individual +1 sites and replicate. A paired Wilcoxon signed-rank test was applied on site-averaged ΔH3 values to confirm the difference between conditions.**(E)** Differential pause release index (PRI), as calculated in **C**, for ERK-induced genes, sorted by log2 fold change (FC) (2h/0h) of PRI in CHD4-depleted cells. Genes on the left have a higher PRI log2 FC (2h/0h) in IAA-treated cells compared to the control and represent the strongest release defect upon CHD4 loss.

**Supplementary Figure 8.**
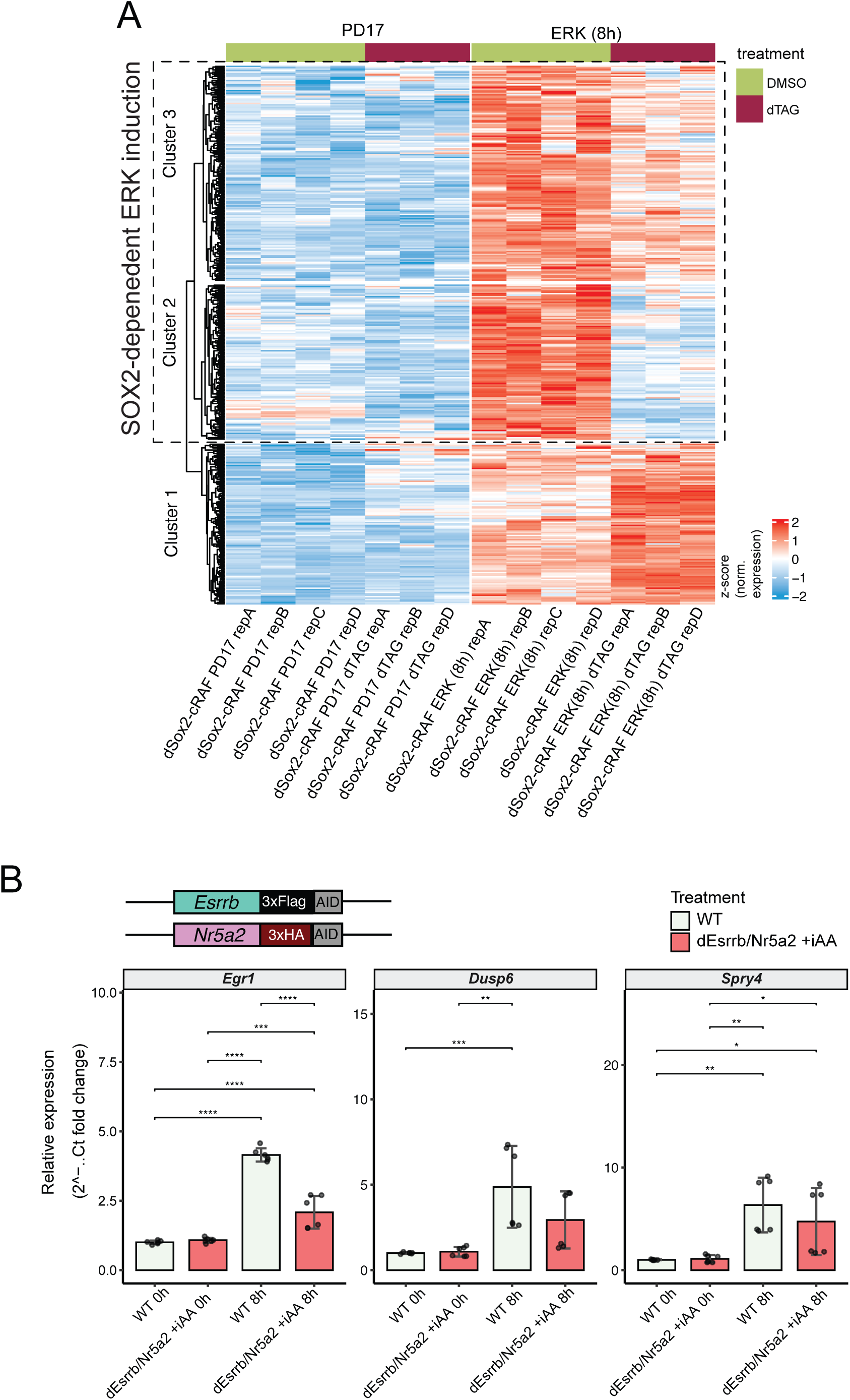
Tethering-mediated recruitment of the pioneers TFs is essential for the activation of ERK-induced genes. **(A)** Heatmap representing z-scored rlog normalized reads of the canonical ERK-induced genes at 8h as determined by DeSEQ analysis (DMSO_PD17 vs DMSO_ERK(8h), padj>0,05 and log2fc >0) in dSOX2-cRaf cells. Each replicate, A, B, C or D, represent different cell passages. Cells were cultured in PD17 for 24h, dTAG13 or DMSO was added 1 h prior to inducing ERK for 8h by adding 4OHT. K-means clustering (k=3) calculated through ComplexHeatmap.**(B)** Representation of *Esrrb* and *Nr25a* alleles targeted in the ERK-inducible cell line to generate to generate the dESRRB/NR25A cell line that allows for the quick degradation of both proteins and the induced activation of ERK signaling with addition of 4OHT. The plots show the relative expression of ERK-induced genes (*Egr1*, *Dups6* and *Spry4*) measured by quantitative PCR (qPCR), in the presence and absence (iAA) of ESRRB and NR52A.

**Supplementary Table 1:** MED24 peaks affected by 2h of ERK induction

**Supplementary Table 2:** Data from the MED24 RIME

**Supplementary Table 3:** Nuclear Proteome of MED24-FLAG cells with ERK induction

**Supplementary Table 4:** CHD4 binding and comparison to SOX2

**Supplementary Table 5:** SOX2 differential binding and motif analysis

**Supplementary Table 6:** Differential ATAC-seq peaks

**Supplementary Table 7:** Canonical ERK-induced and repressed genes at 2h

**Supplementary Table 8:** PRI fold changes and ERK induction in the absence of SOX2

